# Kidney organoid reproducibility across multiple human iPSC lines and diminished off target cells after transplantation revealed by single cell transcriptomics

**DOI:** 10.1101/516807

**Authors:** Ayshwarya Subramanian, Eriene-Heidi Sidhom, Maheswarareddy Emani, Nareh Sahakian, Katherine Vernon, Yiming Zhou, Maria Kost-Alimova, Astrid Weins, Michal Slyper, Julia Waldman, Danielle Dionne, Lan T Nguyen, Jamie Marshall, Orit Rosenblatt-Rosen, Aviv Regev, Anna Greka

## Abstract

Human iPSC-derived kidney organoids have the potential to revolutionize discovery, but assessing their consistency and reproducibility across iPSC lines, and reducing the generation of off-target cells remain an open challenge. Here, we used single cell RNA-Seq (scRNA-Seq) to profile 415,775 cells to show that organoid composition and development are comparable to human fetal and adult kidneys. Although cell classes were largely reproducible across iPSC lines, time points, protocols, and replicates, cell proportions were variable between different iPSC lines. Off-target cell proportions were the most variable. Prolonged *in vitro* culture did not alter cell types, but organoid transplantation under the mouse kidney capsule diminished off-target cells. Our work shows how scRNA-seq can help score organoids for reproducibility, faithfulness and quality, that kidney organoids derived from different iPSC lines are comparable surrogates for human kidney, and that transplantation enhances their formation by diminishing off-target cells.

## Introduction

Kidney diseases affect ∼800 million people worldwide(Coresh et al., 2007). Despite the enormous disease burden, therapeutic innovation has lagged(Inrig et al., 2014), owing in part to the lack of appropriate models that reflect the cellular complexity of the human kidney. Technologies to generate kidney organoids from human induced pluripotent stem cells (iPSC)(Cruz et al., 2017a; Forbes et al., 2018a; Freedman et al., 2015; Morizane and Bonventre, 2017a; Morizane et al., 2015; Przepiorski et al., 2018; Taguchi and Nishinakamura, 2017; Taguchi et al., 2014; Takasato et al., 2015a, 2016a; Tanigawa et al., 2018a) provide a promising avenue to further help advance our understanding of disease mechanisms and expedite therapeutic development.

To harness the full potential of iPSC derived kidney organoid technology, we must address critical unsolved questions about their reproducibility, faithfulness, and quality. First, we must establish organoid reproducibility: the comparability and range of variability in cellular composition and state between different iPSC lines from normal individuals across replicates and protocols (building on previous efforts to draw comparisons between iPSC and embryonic stem cell (ESC) derived organoids(Wu et al., 2018) or bulk RNASeq data comparing iPSC derived organoids(Phipson et al., 2019)). This is critically important, because individual patient iPSCs offer significant advantages over ESCs for precision medicine and drug development projects(Boreström et al., 2018; Burrows et al., 2016; Takasato et al., 2016b). Second, we must define their faithfulness: how well organoids across many iPSC lines recapitulate kidney development and disease-associated genes at single cell resolution. Third, we need to define their quality. Since off-target cells interfere with organoid quality, we must understand how to drive their removal to more faithfully reproduce the human kidney(Wu et al., 2018).

Here, we address these critical questions by combining scRNA-Seq analysis of cellular composition with immunofluorescence validation to build a comprehensive atlas of human kidney organoids across iPSC lines, replicates, differentiation protocols, and developmental time, and after organoid transplantation into mouse.

## Results

### A ∼400K single cell census from 47 kidney organoid states derived from 4 human iPSC lines

To compare organoids at single cell resolution, we generated them according to two protocols, from each of four different human iPSC lines generated by different methods (episomal or Sendai virus; **Supplementary Fig. 1A**), and at multiple time points along differentiation, and profiled them by droplet-based scRNA-seq (**Fig. 1A**). First, we used the ML protocol(Takasato et al., 2016b) and generated organoids from each of two commercial lines (Thermo Fisher (ThF, female) and Alstem (AS, male)), and two obtained from human healthy donors (N1, female and N2, male) (**Supplementary Fig. 1A**). We profiled single cells at the day 0 (D0) iPSC state, at two critical milestones, day 7 (D7; immediately prior to cells being plated for 3D self-organization and nephrogenesis), and day 15 (D15; when growth factor treatment ends), and finally, at day 29 (D29) when the organoids are mature. This allowed us to assess the reproducibility within a protocol and the impact of cell-line-specific differences in iPSC pluripotency and/or subsequent differentiation. Second, we generated kidney organoids using the JB protocol(Morizane and Bonventre, 2017b) from ThF iPSCs at D29 (**Fig. 1A**). To account for technical variability, we sequenced single cells from three different iPSC replicates (D15, D29, all lines) as well as an additional, independent passage (AS line at D7, D15, D29 and TF at D7). We successfully profiled 382,465 single cells from organoids across all four iPSC lines (**Fig. 1B** and **Supplementary Fig. 1B**) and compared them to 3,417 cells profiled from the unaffected kidney portion of a tumor nephrectomy from an adult male (with no known kidney diseases) (**Fig. 1C**). We determined the cellular composition of organoids using unsupervised graph-based clustering followed by *post-hoc* annotation (**Methods**) with signatures of cell types (**Supplementary Fig. S2**, **Supplementary Table 1**) and cell cycle genes(Kowalczyk et al., 2015).

**Figure. 1.**
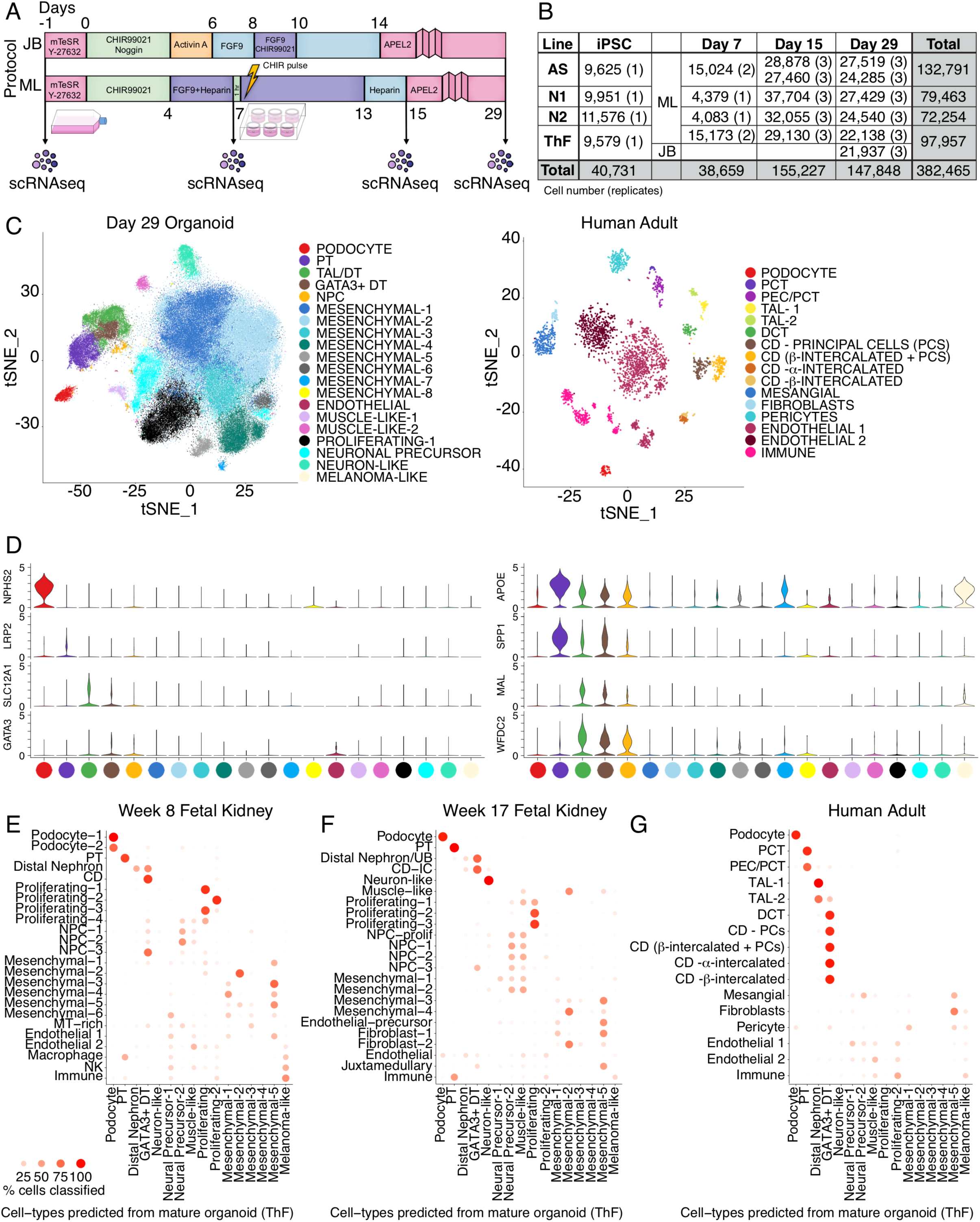
Mature kidney organoids from four different human iPSC lines contain most major nephron cell classes. (A) Schematic of differentiation protocols for kidney organoids derived from human iPSCs by the Bonventre (JB) and Little (ML) protocols. Single-cell sequencing time-points as shown. (B) Table summarizing single cells profiled across four different human iPSC cell lines (AS, N1, N2, ThF) using two different protocols (ML, JB) across four time points (iPSC, Day 7 (D7), 15 (D15) and 29 (D29)). Replicates are indicated in parentheses. (C) t-SNE plot of composite single cell transcriptomic profiles from all 4 iPSC D29 kidney organoids (left) and human adult kidney (right). Cluster color annotations as shown. (D) (left) Violin plots showing expression of canonical adult human kidney markers used to identify nephron clusters in D29 organoids: podocyte (NPHS2), proximal tubule (PT; LRP2), thick ascending limb/distal nephron (TAL; SLC12A1) and immature distal nephron (GATA3+ DT). Mesenchymal and off-target cells were also detected. (right) Violin plots of data-derived tubular markers revealed APOE proximal to distal gradient of expression, proximal marker SPP1, and distal markers WFDC2 and MAL. (E) Random forest classifier trained on D29 ML_ThF organoid gene expression profiles accurately predicted respective nephron cell classes in human first and second trimester fetal kidney (E, F) as compared to adult kidney (G). The Y-axis labels in each panel represents the assigned cell class in each human sample. The X-axis labels are cell classes at D29 (ML_ThF organoids). The size and intensity of the red dot represents the % of cells in the Y-axis cell-types classified as the corresponding X-axis cell-class, e.g., all human adult podocytes are correctly classified as podocytes, while distal convoluted tubule (DCT) is predicted to be most similar to the GATA3+ DT organoid cell class.

### Mature organoids from all iPSC lines contain predominant nephron cell classes

D29 organoids from all 4 iPSC lines contained cells representative of segments of a developing nephron (**Fig. 1C, D** and **Supplementary Fig. 2)**: podocytes (NPHS2, podocin; NPHS1, nephrin; SYNPO, synaptopodin; WT1, Wilms Tumor 1), proximal tubular (PT) cells (LRP2, megalin), thick ascending limb (TAL; SLC12A1, Na-K-Cl cotransporter), and distal nephron cells (CDH1, E-cadherin; AQP2, aquaporin 2; GATA3). There was no cluster enriched for SLC12A3 (Na-Cl symporter; **Supplementary Fig. 2**), a canonical marker of the distal convoluted tubule (DCT). The organoid single cell profiles retained the proximal (podocyte) to distal axis of the human nephron (**Fig. 1C**, left) on visualization of the data using t-distributed Stochastic Nonlinear Embedding (tSNE), unlike the discrete clusters seen in adult kidney (**Fig. 1C**, right). We identified data-derived markers (**Supplementary Table 2**), including osteopontin (SPP1)(Xie et al., 2001) in the proximal tubular cell cluster, and WFDC2(LeBleu et al., 2013) and MAL(Carmosino et al., 2010) in the distal nephron cluster (**Fig. 1D** and **Supplementary Fig. 3A**), in line with single cell RNA-Seq studies of human adult and fetal kidney(Lindström et al., 2018a). In all 4 iPSC lines, we observed a novel proximal to distal tubular gradient of APOE (a gene associated with diabetic kidney disease(Araki, 2014))(**Fig. 1D**). Thus, D29 organoids reproducibly developed podocytes, proximal tubular cells, and cells consistent with the loop of Henle and distal nephron (but without a defined distal convoluted tubule (DCT) or collecting duct (CD) segment as seen in adult kidney).

D29 organoids also contained Nephron Progenitor Cells (NPC) enriched in PAX2, LHX1 and PAX8 (top cluster-specific differentially expressed (DE) genes). The majority of the organoid single cells (70% on average) were mesenchymal (**Fig. 1C**), grouped in 8 subsets (Mesenchymal 1-8) enriched for markers of progenitor and differentiating cell types (**Fig. 1C; Supplementary Fig. 2**). Prominent non-kidney off-target populations(Wu et al., 2018), absent in adult human kidney (**Fig. 1C, Supplementary Fig. 2**), were found in D29 organoids, including melanoma-like cells (PMEL), SOX2-positive(+) neuronal precursors, STMN2+ neuron-like cells, and MYOG+ muscle-like cells, as reported previously(Phipson et al., 2019; Wu et al., 2018). A rare population of endothelial cells was also observed (**Fig. 1C** and **Supplementary Fig. 2**).

### Mature organoids most closely resemble fetal kidney

We related the cell clusters from D29 human kidney organoids (ThF line) to human tissue by comparing them to fetal kidney from the first (8 weeks) and second (17 weeks) trimesters(Lindström et al., 2018b; Young et al., 2018), and to adult kidney (**Fig. 1C**, right). We used a classification based approach (random forest classifier)(Pandey et al., 2018) to assess the relation between each cell cluster in these tissues to each organoid cluster (**Fig. 1E-G, Methods**).

Overall, cells were most similar to those from first and second trimester fetal kidneys, largely consistent with previous studies using bulk RNA-Seq data(Takasato et al., 2015b). All nephron lineages were accurately classified by the algorithm (**Fig. 1E-G**). Compared to those in adult human kidney (**Fig. 1C**), all organoids contained cells of human nephron segments, but with poorly differentiated distal tubular cells. In contrast, in the adult kidney, we identified distinct proximal, loop of Henle, DCT and CD sub-clusters, including principal cells (PC) and α- and β-intercalated cells (IC), as well as endothelial and immune cells (**Fig. 1C**, **Supplementary Fig. 3B**).

Interestingly, some of the mesenchymal cell types in kidney organoids were predictive of cell types in fetal and adult kidney, suggesting that these are on-target mesenchymal cells. Fetal kidney cell types (mesenchymal, endothelial, and fibroblasts) were classified to mesenchymal-1, -2, and -5 clusters in organoids (**Fig. 1E, F, Supplementary Table 2**), while mesangial cells, fibroblasts, and pericytes in adult kidney were classified to mesenchymal-1 and -5 in organoids (**Fig. 1G**). Of interest, mesenchymal-2 cells in organoids were enriched for MGP, GAS2, PRXX2 and FOXP2(Wang et al., 2018), known markers of fetal mesangial cells in the developing kidney. Similarly, mesenchymal-5 cells in organoids were enriched for fetal stromal genes SULTIE1, DKK1(Iglesias et al., 2007), NR2F1(Schwab et al., 2003), ID3, ZEB2, COL6A3(Menon et al., 2018) and DCN. In contrast, mesenchymal types that did not map to human kidney cell types included genes associated with cartilage formation(Ohba et al., 2015) (ACAN, SOX9, MATN4, LECT1, EPYC, COL9A3 and COL9A1)(**Fig. 1E, F**).

We validated cell type markers using immunofluorescence (IF) staining (**Fig. 2A-C**, **Supplementary Fig. 4, 5A**). Markers for podocyte (WT1), proximal tubule (LTL) and distal nephron (CDH1) were detected across all lines and protocols, although expression of GATA3, a marker of the developing ureteric bud, could only be detected in N2 and ThF(Lee et al., 2015; Takasato et al., 2016b) (**Fig. 2C**). We also validated the markers used to annotate specific cell clusters: NPHS1(Greka and Mundel, 2012) (podocyte cluster), co-localized with the podocyte-specific marker synaptopodin (SYNPO)(Mundel et al., 1997) (**Fig. 2D**); LRP2(Christensen and Birn, 2002) (proximal tubule cluster), co-localized with LTL, a lectin specific to the proximal tubule (**Fig. 2E**); CDH1(Prozialeck et al., 2004) (distal tubular compartment) marked tubules distinctly from proximal LRP2/LTL positive tubular structures (**Fig. 2E**). We also validated MEIS1, a gene enriched in kidney mesenchymal cells, by IF staining, where it was shown to localize appropriately to the interstitium, defined as the cellular space outside laminin positive basement membrane structures (LAMA1) (**Fig. 2F**).

**Figure 2.**
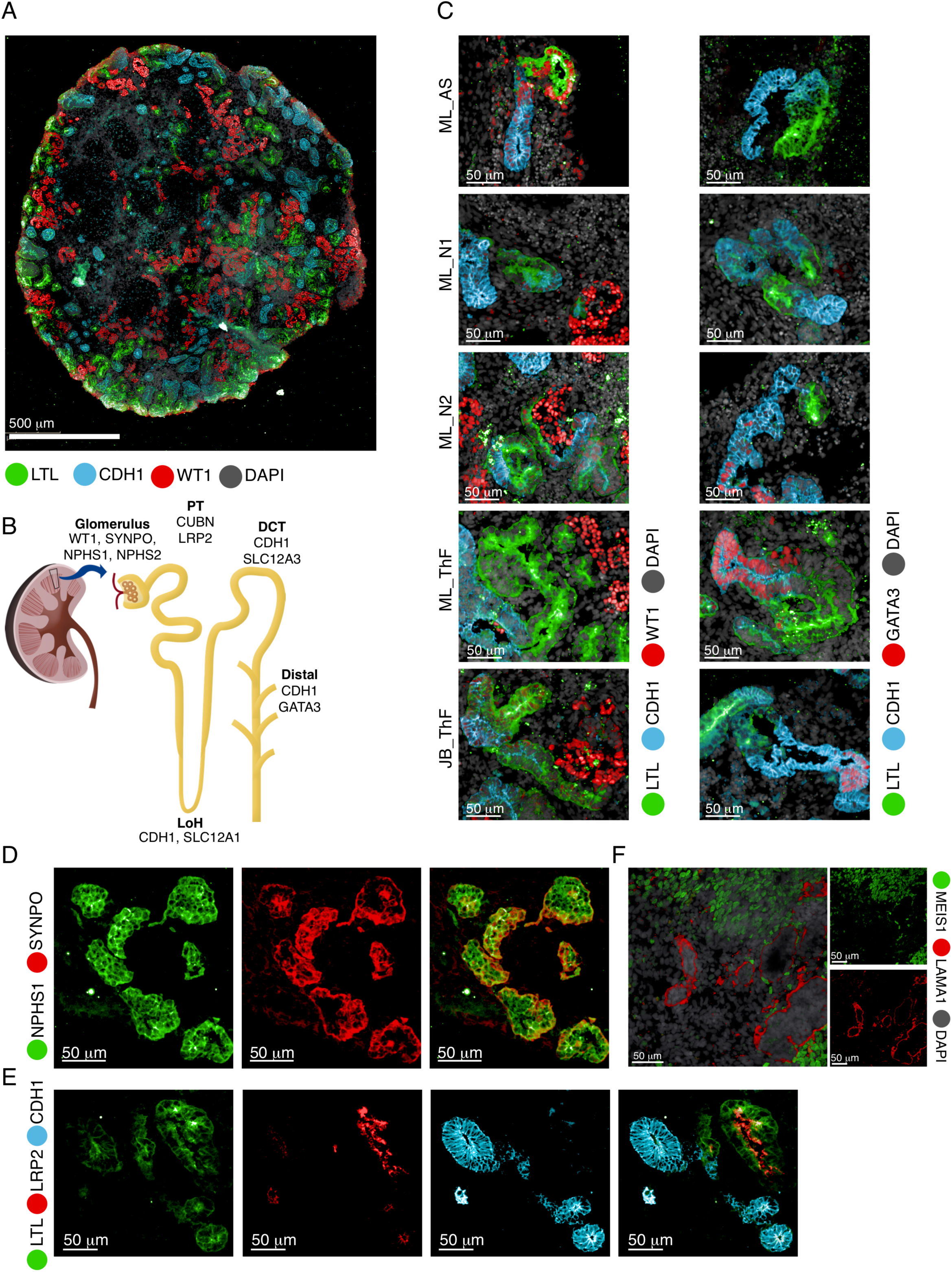
IF validation of markers derived from the single cell data in mature kidney organoids. (A) IF staining of an entire kidney organoid with segment specific markers as shown. (B) Schematic of kidney nephron with major cell types and canonical markers annotated. (C) Immunofluorescence staining of D29 kidney organoids for podocyte (WT1), proximal tubule (LTL), and distal tubule (CDH1 and GATA3) across two protocols (JB, ML) and four cell lines (AS, N1, N2, ThF). IF staining for validation of markers identified in the single cell data: (D) NPHS1 co-localized with the podocyte-specific marker SYNPO and (E) LRP2 co-localized with the proximal tubular marker LTL (bottom). (F) IF staining validation for MEIS1-positive mesenchymal cells in D29 organoids. LAMA1 indicates basement membranes. MEIS1 staining of mesenchymal cells appropriately surrounds LAMA1-defined tubular nephron structures.

### Cell proportions vary depending on iPSC line

Cell type proportions in mature D29 organoids were consistent between replicates of independent clones, but they varied between iPSC lines (**Fig. 3A, B**). We quantified the differences in cell proportions by computing the Jenson-Shannon Divergence(Yi-Rong Peng, Karthik Shekhar, Wenjun Yan, Dustin Herrmann, Anna Sappington, Greg S. Bryman, Tavé van Zyl, Michael Tri. H. Do, 2018) (JSD, a measure of compositional difference between 2 frequency vectors with a value between 0 and 1, where smaller values mean more similar) within- and between-iPSC lines (**Fig. 3C**). Differences between lines (average JSD×0.18, sd × 0.13) superseded differences between independent clones within the same line (average JSD 0.01, sd 0.01), and between different protocols for the same line (average JSD× 0.06, sd×0.02) (Fig. 3C). N2 was most different (divergent) from both AS and N1 iPSCs that were the most similar. For example, podocytes were captured across all experimental conditions (**Fig. 3A** and **Supplementary Fig. 5B**), but varied between 1.33% (N1, averaged across replicates) to 2.58% (ThF) between lines on the ML protocol. AS, passage 1 (AS-1) was an outlier, with lower overall nephron numbers (0.3%). The JB protocol captured an average of 0.81% podocytes. Similarly, the distal nephron compartment (including TAL and GATA3+ distal nephron cells) ranged in average proportion from 2.34% (N1) to 6.95% (ThF), with an average 1.45-fold-change between protocols. In general, N1 organoids had the lowest average nephron cell proportions, followed by AS, N2 and ThF. GATA3+ distal-like cells were more abundant in ThF iPSCs, independent of protocol; GATA3 expression in AS and N1 was lower than in ThF and N2 (**Supplementary Fig. 2**), as confirmed by IF (**Fig. 2C**).

**Figure 3.**
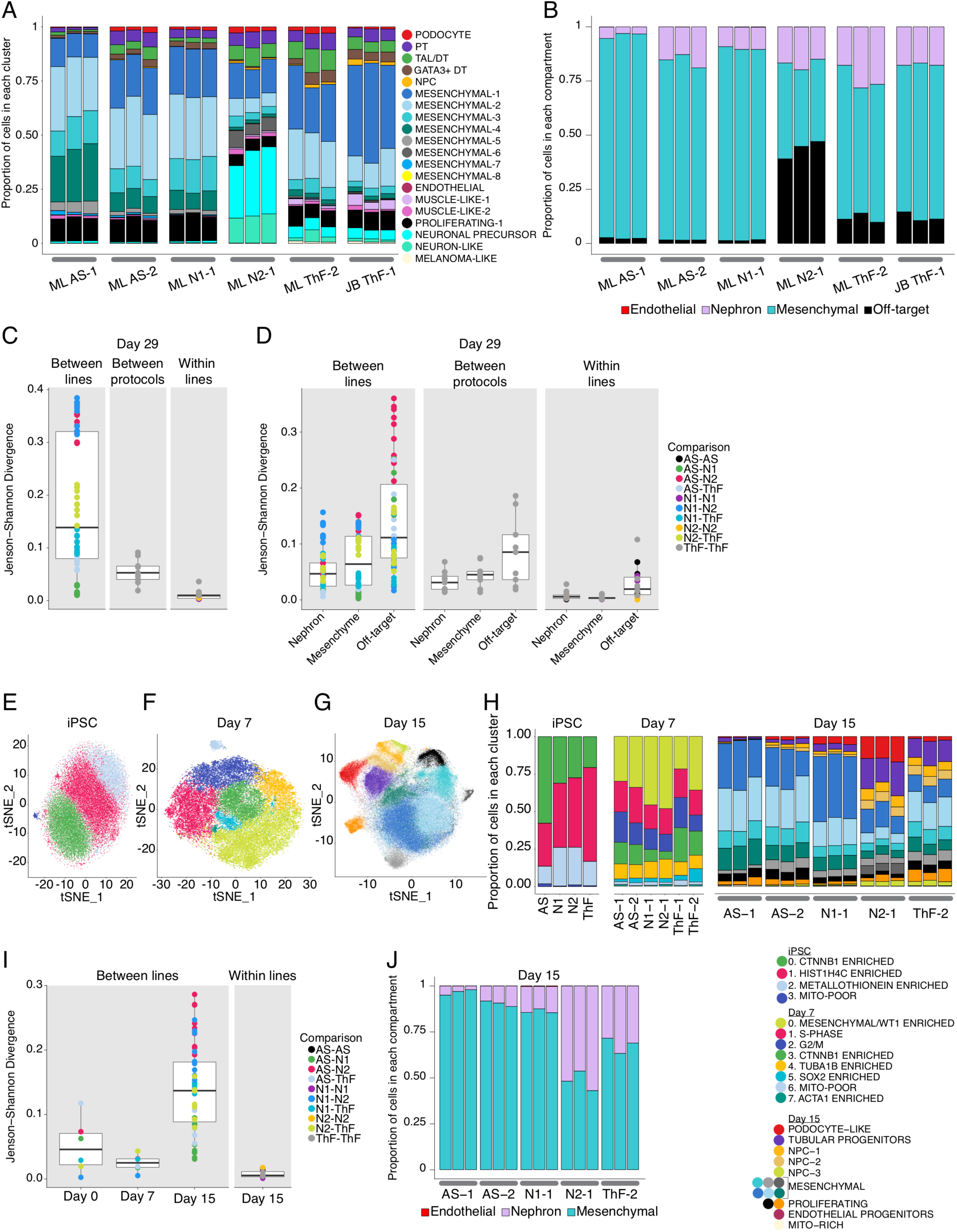
Variability in cell type proportions detected by scRNA-Seq at D15. (A, B) Relative proportions of endothelial, nephron, mesenchymal and off-target cell clusters across all replicates of D29 organoids. Annotations as shown. (C, D) Comparison of cell-type composition between D29 organoids as determined by boxplots of the Jenson-Shannon Divergence (JSD) method. Each point on the plot is a pair-wise (color) measure of JSD between 2 organoids. Legend indicates annotation for pairs of iPSC lines. (C) Organoid compositional differences are greater between lines than between different protocols for the same line or between replicates of the same line and protocol (within lines). (D) Organoid compositional heterogeneity is greatest in the off-target compartment followed by the mesenchyme and the nephron compartment in all three comparison groups (between lines, between protocols and within lines). t-SNE plot of single cells from (E) iPSC, (F) D7, and (G) D15 of the organoid differentiation protocol. (H) Comparison of relative cell type proportion across iPSC lines of cell clusters shown in (E-G). (I) Compartment specific cell-type proportions at D15 across all lines and replicates. (J) Compositional differences are greatest between lines at D15 and the least at D7.

To make higher level comparisons, we looked at 4 groups: nephron, mesenchymal, off-target and endothelial compartments (**Fig. 3B, Supplementary Table 3**). The nephron compartment was on average 16.7%, of all cells (9.89% in N1 to 24.1% in ThF). The N1 line had the largest average relative proportion of endothelial cells (0.1%), with a global average of 0.05% across all lines. Mesenchymal cells were 82.8% of all cells in AS and 88.6% in N1 organoids, but only 64.2% in ThF and 39.2% in N2. Off-target cells varied markedly by iPSC line, from 1.6% in AS-1 and N1 to 43.7% in N2 (**Fig. 3B**). To determine variability within each compartment, we again computed the JSD (**Fig. 3D**). The nephron and mesenchymal compartments were consistent (nephron mean JSD between lines × 0.05, sd × 0.038; mesenchymal mean JSD × 0.07, sd × 0.05), whereas the off-target compartment was variable between lines (mean JSD × 0.14, SD × 0.09) and protocols (**Fig. 3D**). N2 organoids were most divergent from AS in off-target composition. Notably, the ratio of mesenchymal to nephron cells was inversely related (Spearman correlation ρ × −0.84) to the proportion of off-target cells. Higher proportion of off-targets (N2, ThF:11.7%, JB ThF: 12.2%) resulted in lower mesenchymal:nephron ratios (N2: 2.29, ThF: 2.66, JB ThF: 4.05) and vice-versa (Off-targets – N1, AS:1.59%; mesenchymal: nephron – N1: 8.96, AS: 5.31). In contrast, the mesenchyme proportions were lower in both adult and fetal kidney, both overall (15.7% in adult, 19.7% in fetal week 17) and relative to nephron cells (0.65 in adult, 0.47 in fetal). In summary, we noticed greater organoid heterogeneity between iPSC lines than between replicates within a line, or between protocols, with the off-target compartment contributing the most variability.

### Variability in cell type proportions detected by scRNA Seq in D15 organoids

To explore whether the iPSC-line associated differences were associated with their basal state(Burrows et al., 2016; Féraud et al., 2016) at D0 or the process of differentiation, we compared the single cell profiles collected at D0,7, and 15 for each organoid culture. Analysis of 42,433 single cells from the 4 undifferentiated iPSC lines at D0 (**Fig. 3E, H** and **Supplementary Fig. 6A**), showed they were comparably pluripotent. To determine if there were clusters of cells primed towards a particular developmental germ layer, we scored organoids using known transcriptional signatures for the three germ layers. We did not observe sub-clusters with signatures for any specific germ layer(Tsankov et al., 2015), or primed for differentiation(Nguyen et al., 2010) (**Supplementary Fig. 7A**). The four cell clusters had representation from each of the iPSC lines (avg JSD × 0.05, sd × 0.04, **Fig. 3H, I**), and from all cell-cycle phases (**Supplementary Fig. 7B**).

Similarly, at D7, developing organoids from all iPSC lines expressed appropriate markers of mesodermal differentiation with actively proliferating cells(Kowalczyk et al., 2015) (**Supplementary Fig. 7C, D**) and little variability between lines (**Fig. 3F, H**). Differences in proportions were small (avg JSD×0.02, sd×0.01, **Fig. 3I**). Notably, clusters 3, 5 and 0 appropriately expressed CTNNB1 and SOX11, SOX2, and WT1, respectively, genes upregulated during the differentiation of intermediate mesoderm (IM) into the metanephric mesenchyme (MM) or the ureteric bud (UB)(Kreidberg et al., 1993; Little and McMahon, 2012; Little et al., 2016) (**Supplementary Fig. 6B**). We did not observe nephron cell types in D7 organoids.

At D15, organoids had multiple subsets of nephron progenitor cells (NPC) and a distinct population of epithelial cells broadly expressing tubular progenitor markers (PAX2, LHX1; **Supplementary Fig. 8A**). Across all iPSC lines, we observed early patterning of the nephron (reflected by expression of proximal (CUBN) and distal (MAL, WFDC2, POU3F3) markers; **Supplementary Fig. 8A**), proliferating cell populations (*e.g.*, cells in the Mesenchymal-4 cluster expressed CTNNB1, PTPRS, CDC42, and SOX11, similar to CTNNB1-expressing cells in clusters in the iPSCs and D7 organoids; **Supplementary Fig. 8A**), and a distinct subset of podocyte-like cells (**Fig. 3G, H; Supplementary Fig. 8A**). In contrast to D7 and D0 (**Fig. 3I**), there were notable differences in composition between iPSC lines (avg JSD × 0.14, sd × 0.07; **Fig. 3G, H, I**). Heterogeneity in endothelial progenitors was consistent with D29: N1 had the highest average relative proportions (0.32% vs 0.05% among other lines; **Fig. 3J**). The proportion of nephron-like cells ranged from 0.08% in AS to 56.9% in N2 D15 organoids (**Fig. 3H, J**). Although, no distinct off-target cells were found at D15, N2 and ThF organoids had a small population of cells expressing the neuronal progenitor SOX2 within the mesenchymal compartment (**Supplementary Fig. 8B**). The mesenchyme:nephron ratios were also lower in N2 and ThF D15 organoids (**Fig. 3J**). Hence, organoids with higher nephron proportions at D15 had a distinct pool of SOX2+ cells, and went on to develop a higher proportion of off-target cells at D29, associated in turn with lower D29 mesenchyme:nephron ratios (**Fig. 3B, J**). Taken together, variability in organoid differentiation was evident by D15.

### Concordant expression of developmental programs across organoids from 4 human iPSC lines

Next, we tested whether known transcription factors (TFs) and critical genes involved in nephrogenesis(Little and McMahon, 2012; Little et al., 2016) are appropriately expressed in the respective clusters in organoids from 4 different iPSC lines, compared to adult kidney (**Fig. 4A, Supplementary Fig 9,10**). Overall, key developmental programs were expressed at expected time points and transitions, in a comparable manner across different iPSC-derived organoids (**Fig. 4A**, **Supplementary Fig. 9,10**). First, core pluripotency TFs were expressed during the iPSC stage and decreased subsequently (D7-D29). Notably, SOX2, a neuronal progenitor marker(Graham et al., 2003), re-appeared in D29 organoids in off-target neuron cells (**Fig. 4A**). Next, at D7, prior to the CHIR pulse (**Fig. 1A**), cells across all clusters had high expression of markers of mesoderm development, such as HAND1(Poh et al., 2014) and HOX11 genes(Wellik et al., 2002). With organoid self-assembly between D7 and D15, IM inducing genes from D7 were downregulated in D15, while nephron progenitor genes were upregulated in a nephron-specific pattern: JAG1, LHX1, PAX2, PAX8 in the epithelial clusters and WT1 in the nephron progenitor population and podocyte-like cells (**Fig. 4A**). At D15 we first observed a SOX17-positive endothelial progenitor cell population that persisted in D29 (**Fig. 4A, Supplementary Fig. 9C**). Nephron patterning became more evident at D29: FOXC2 and WT1 were prominent in podocytes, whereas IRX3 and POU3F3 had a distinct distal pattern (**Fig. 4A**, **Supplementary Fig. 10**). Developmental programs were comparable across all iPSC lines (**Supplementary Fig. 9, 10**).

**Figure 4.**
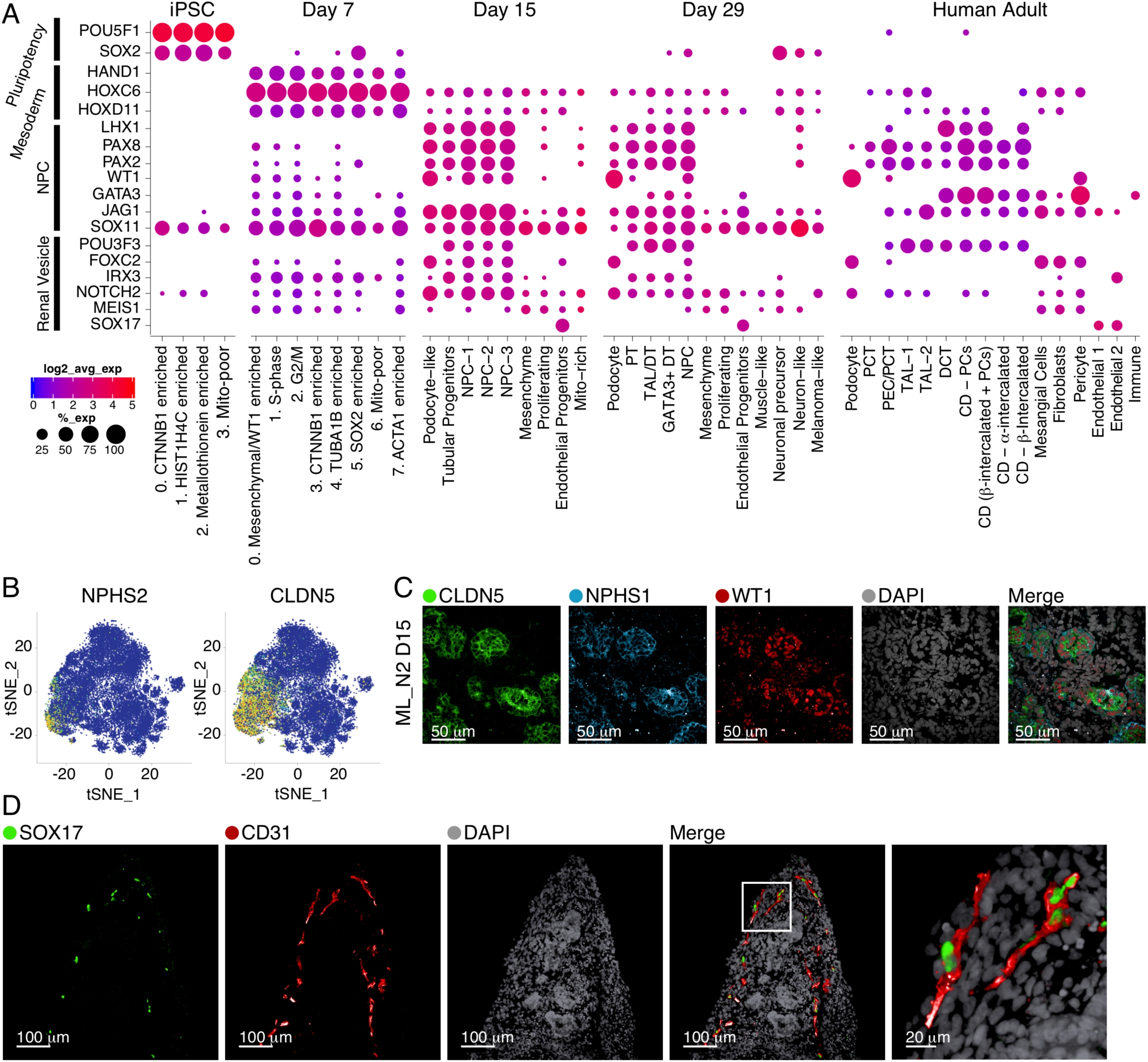
Concordant expression of developmental programs across organoids from 4 human iPSC lines. (A) Dot plot comparison of expression patterns for major nephrogenesis markers across organoid differentiation time points (iPSC, D7, D15, and D29, averaged across four cell lines, ML protocol) and human adult kidney. The size of the dot represents the proportion of cells in the cluster that express the gene. Dots are shown only when the gene is expressed in at least 5% of the cells in the respective cluster. (B) Canonical (NPHS2) and data-derived (CLDN5) podocyte marker genes superimposed in tSNE plots from D15 organoids (N2 line, ML protocol). (C) IF staining of D15 kidney organoid (N2 line, ML protocol) for CLDN5 as a marker of early podocyte differentiation derived from the single cell data. Additional canonical podocyte markers (NPHS1, WT1) and DAPI staining as shown. (D) IF staining of D29 kidney organoid (AS line, ML protocol) for SOX17 and CD31, markers of endothelial cells.

Of several data-derived markers enriched in the podocyte population at D15 (CTGF, CLDN5, SOST, SPARC; **Fig. 4B, Supplementary Fig. 8C**), we validated CLDN5 (claudin 5, an integral membrane protein controlling tight junctions(Yuan et al., 2012) in many cells including podocytes(He et al., 2007)) in nephrin (NPHS1) and WT1-positive podocytes of D15 organoids (**Fig. 4C**). To validate the endothelial precursors identified in our analysis of D15 organoids, we stained for SOX17 (localized to endothelial cell nuclei) and showed that it co-localized with CD31, a classical endothelial marker (**Fig. 4D**).

### Genes associated with kidney diseases are expressed in expected compartments across organoids from 4 human iPSC lines

To assess the applicability of kidney organoids from different iPSC lines for the study of genetic kidney diseases(Cruz et al., 2017b; Forbes et al., 2018b; Tanigawa et al., 2018a), we determined the expression of genes causing congenital anomalies of the kidney and urinary tract (CAKUT), hereditary renal cystic (HRC) diseases and hereditary tumor syndromes(Brown et al., 2014; Hildebrandt, 2010; Vivante and Hildebrandt, 2016; Warejko et al., 2018) (**Fig. 5A,B, Supplementary Fig. 11,12**), as well as genes from Genome Wide Association Studies (GWAS) of chronic kidney disease (CKD)(Pattaro et al., 2016) (**Supplementary Fig. 13**) and mendelian glomerular diseases(Ashraf et al., 2018) (**Supplementary Fig. 14**).

**Figure 5.**
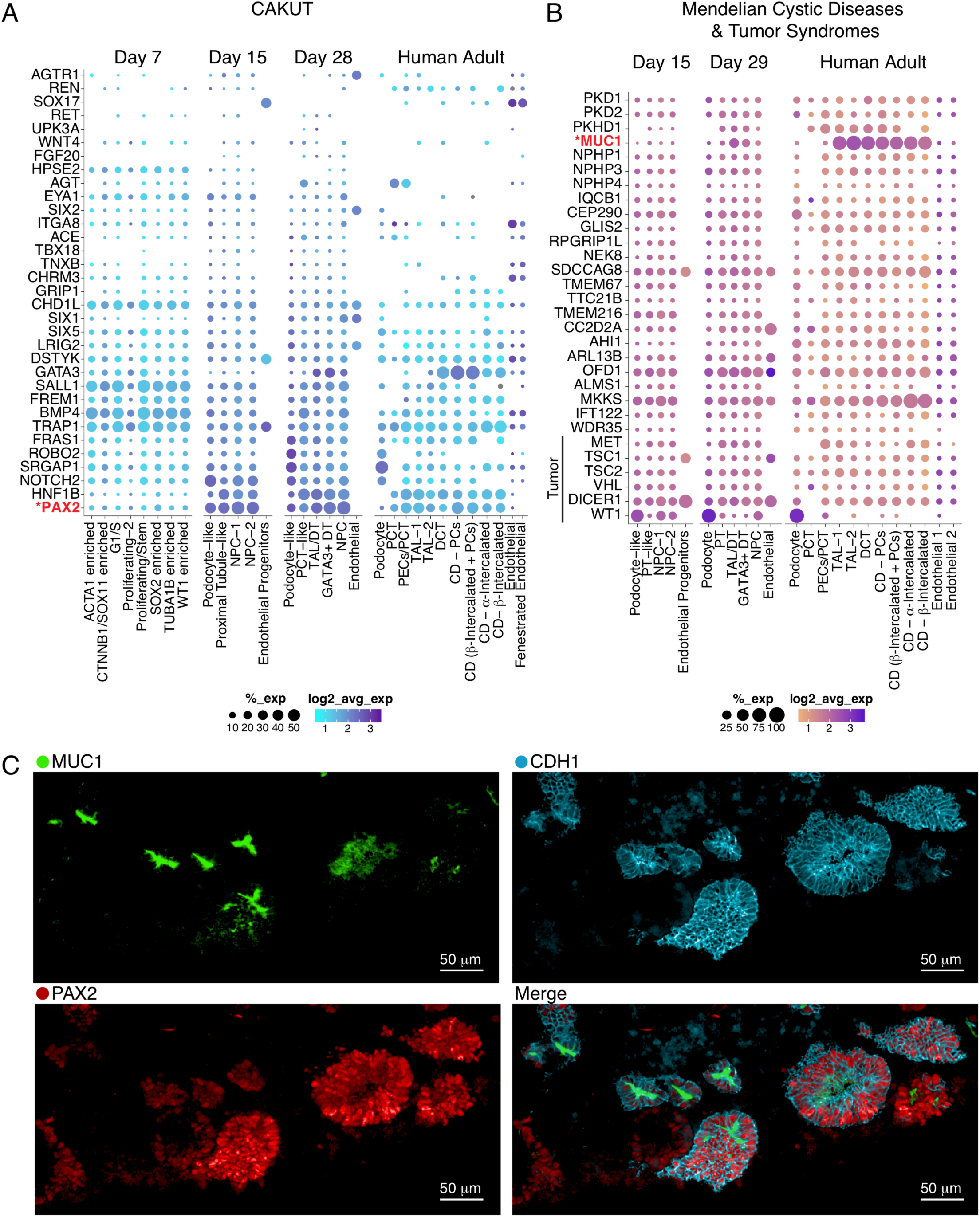
Genes associated with kidney diseases are expressed in expected compartments across organoids from 4 human iPSC lines. Dot plot comparison of cell type-specific gene expression (ThF line, ML protocol) in organoids and human adult kidney. Genes implicated in monogenic causes of (A) congenital abnormalities of the kidney and urinary tract (CAKUT), and (B) mendelian renal cystic diseases and tumor syndromes. (C) IF staining validates the expression of MUC1 and PAX2 in the distal tubule in D29 kidney organoids. CDH1 serves a distal tubular marker. Note appropriate apical expression of MUC1 in distal tubular epithelial cells.

Organoids from all lines reproducibly expressed the majority of genes associated with progressive kidney diseases (including adult onset diseases) in at least one compartment. Genes associated with embryonic and early childhood abnormalities (CAKUT and HRC) were broadly expressed in developing organoids from all iPSC lines as early as D7 (i.e. CHD1L(Brockschmidt et al., 2012) at D7). Appropriately, developmental genes enriched in organoids were absent in adult kidney (i.e. RET, HPSE2). Some genes achieved high levels of expression and cell-type specificity in mature organoids, in a pattern similar to adult kidney. For example, PTPRO (from CKD GWAS and monogenic glomerular diseases) was appropriately expressed in D29 and adult human podocytes(Wharram et al., 2000) (**Supplementary Fig. 13, 14**). Similarly, PAX2 and MUC1 (from CAKUT and cystic diseases, respectively) were highly expressed in D29 distal tubules, similar to their human adult expression pattern(Leroy et al., 2002) (**Fig. 5A, B**; **Supplementary Fig. 11,12**). We validated these markers derived from scRNA-Seq analysis by IF staining, which confirmed the co-expression of PAX2 and MUC1 in CDH1+ distal tubular epithelial cells in mature organoid sections (**Fig. 5C**). In summary, kidney organoids from 4 different iPSC lines could serve as reasonably faithful surrogates of human kidney tissue for the study of a broad array of kidney diseases.

### Organoid transplantation in mice diminishes off-target cells and enhances organoid quality and composition

We hypothesized that the reduction of off-target cells may improve organoid composition and overall quality. First, we determined that prolonged organoid culture *in vitro*, up to 51 days, did not reduce the portion of off-target cells, as assessed by scRNA-seq of organoids grown *in vitro* at day 32 (D32) and day 51 (D51) (**Fig. 6A, B; Supplementary Fig. 15**). D32 and D51 organoids had most populations present in D29 organoids, including muscle, neuronal and melanoma off-target cell clusters. In particular, we noted 2 clusters of neuronal off-target cells at D29, D32 and D51: STMN2+ neuron-like cells (D29, D32) and SOX2+NTRK2+ neuronal-precursor cells (D29, D32, D51; **Supplementary Fig. 2, 15B**).

**Figure 6.**
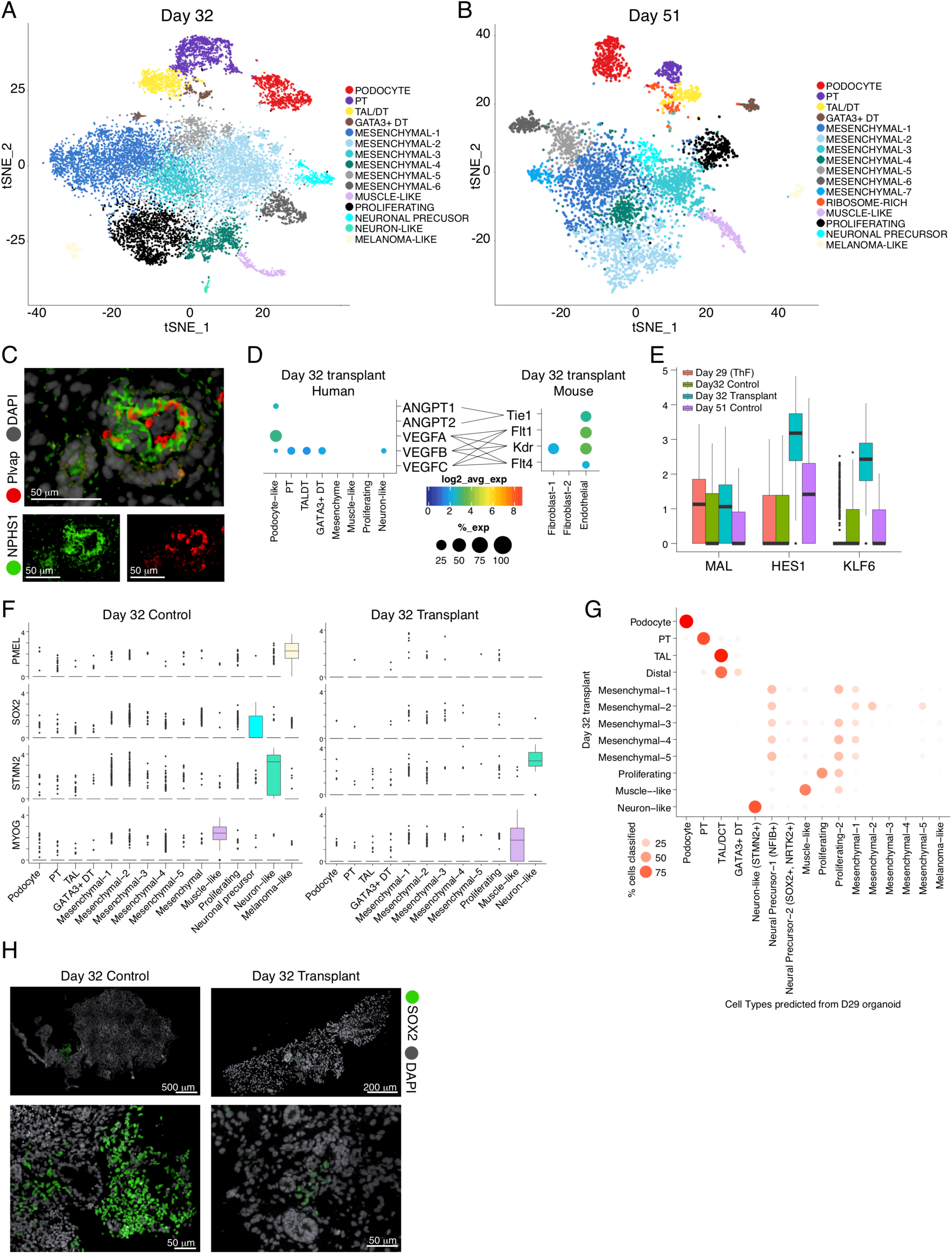
Transplantation of human organoids into mouse diminishes off-target cells and improves organoid quality. t-SNE plots of organoids in prolonged *in vitro* culture reveal cell clusters in (A) D32 and (B) D51 organoids similar to clusters from D29 organoids. (C) Transplanted and vascularized organoid. IF staining shows human podocytes (anti-human NPHS1 antibody) at the outer perimeter and mouse endothelial cells (anti-mouse Plvap antibody) lining the internal perimeter of a glomerular structure. Gray, human and mouse nuclei (DAPI). (D) Dot plots indicating ligand and receptor pairs involved in putative cross-talk between human podocytes and mouse endothelial cells of transplanted organoids. (E) Box plots of expression levels for KLF6 and HES1 in the distal MAL+ cluster from D29, D32 and D51 organoids grown *in vitro*, compared to D32 transplanted organoids. (F) Box plots demonstrate that PMEL+ melanoma cells and SOX2+ neuronal cells were diminished after transplantation. MYOG+ muscle cells and STMN2+ neuron-like cells persisted. (G) Random Forest Classifier shows the relation of cell clusters in transplant organoids compared to organoids grown *in vitro*. Melanoma and SOX2+ neuronal precursor cells are not detected in D32 transplant organoids. (H) IF staining validation for SOX2 (green) in D32 control organoids compared to diminished abundance in D32 transplanted organoids. Human and mouse nuclei (DAPI, gray).

Kidney development requires signaling from adjoining vasculature(Kitamoto et al., 1997), but endothelial cells were minimal in organoids grown *in vitro* (**Fig. 1C and 3A**). Organoid transplantation under the kidney capsule of immunodeficient mice allows mouse endothelial cells to infiltrate the transplanted kidney organoid and promote vascularization(Bantounas et al., 2018; van den Berg et al., 2018; Tanigawa et al., 2018b; Xinaris et al., 2012). We therefore hypothesized that organoid transplantation may improve organoid composition and reduce off-target cells.

To test our hypothesis, we transplanted D18 organoids (ThF line) under the mouse kidney capsule of immunodeficient mice, recovered them 14 days following transplantation (“D32 organoid transplants”), and profiled them by scRNA-Seq, along with D32 control organoids grown *in vitro* (**Supplementary Fig. 16A-D**). We confirmed that transplanted human organoids were vascularized by mouse endothelial cells(Stan et al., 2012): IF showed mouse Plvap-positive endothelial cells apposed to human podocytes (NPHS1) (**Fig. 6C, Supplementary Fig. 16B**). We distinguished the human from mouse cells by read alignment to the combined reference (**Methods**) and confirmed that the human nephron cells in D32 transplants (**Supplementary Fig. 16D, E**) corresponded to D29 nephron clusters, with VEGFA and ANGPT1 most highly expressed in podocytes (**Fig. 6D**). Correspondingly, the VEGF receptors Flt1, Kdr and Flt4(Ferrara, 1999; Kanno et al., 2000; Sharmin et al., 2016; Tufro et al., 1999) and the angiopoietin receptor Tie1 were highly expressed in mouse Plvap-positive endothelial cells (**Fig. 6D**), suggesting that the cell types could interact.

The transplanted organoid epithelial cells had increased expression of genes suggestive of more mature states compared to *in vitro* D32, D29, or D51 controls. Specifically, both KLF6, a TF involved in the development of the ureteric bud and the kidney collecting duct(Fischer et al., 2001), and the Notch effector HES1, which plays a role in proximal to distal patterning in kidney nephrogenesis(Piscione et al., 2004), were upregulated in MAL+ distal cells from D32 transplanted organoids (**Fig. 6E**).

Strikingly, we found little off-target expression of SOX2 (neuronal precursor cells) or PMEL (melanoma cells) in D32 transplanted organoids compared to controls, suggesting that transplantation diminished off-target cells (**Fig. 6F**; **Supplementary Fig. 17A**). MYOG positive muscle cells persisted in D32 transplanted organoids (**Fig. 6F**, **Supplementary Fig. 17A**). We also noted that a rare STMN2+ neuronal cluster persisted in D32 transplants. Interestingly, this cluster was uniquely and highly correlated with the neuronal cluster in week 17 fetal kidney (Spearman ρ × 0.82; **Fig. 1F, Supplementary Fig. 17B**). A set of enriched genes (GAL, CHGA, CHGB(Dressler, 2006); **Supplementary Fig. S18**) was shared between this cluster in fetal kidney and in D32 transplanted organoids (**Supplementary Fig. S18A**). Further analysis revealed that in fact CHGA and CHGB were detectable in a small number of cells within the STMN2+ cluster in D29 control organoids (**Supplementary Fig. S18B**) and transplantation selected for this STMN2+/CHGA+/CHGB+ cluster (**Supplementary Fig. S18A**), suggestive of a more fetal-kidney-like state.

We applied a classifier to further assess the relation of all cell clusters in transplanted organoids to organoids grown *in vitro* (**Fig. 6G**). Consistently, no transplanted organoid cells corresponded to SOX2+NTRK2+ neural precursors or PMEL+ melanoma-like cells (**Fig. 6G**). In line with these findings, IF staining showed diminished SOX2 staining in D32 transplanted organoids compared to numerous SOX2 positive cells in controls (**Fig. 6H**). Taken together, these data showed that transplantation reduced off-target cells, improving organoid maturity and quality.

### Discussion

In this study, we present a comprehensive atlas of human kidney organoids in comparison to human adult and fetal kidneys at single cell resolution spanning multiple iPSC cell lines, time points, protocols, and replicates including transplantation into mice, at average sequencing depths of 10,000 reads per cell in a total of 415,775 cells. This analysis provided answers to several critical questions regarding organoid reproducibility, faithfulness and quality.

First, we compared kidney organoids derived from multiple iPSC lines with single cell resolution. The large number of cells and replicates sequenced across different conditions in this study afforded us the opportunity to retrieve numerous cell types and get robust results from statistics such as the Jenson-Shannon Divergence(Yi-Rong Peng, Karthik Shekhar, Wenjun Yan, Dustin Herrmann, Anna Sappington, Greg S. Bryman, Tavé van Zyl, Michael Tri. H. Do, 2018) to gain quantitative insights into organoid reproducibility. By comparison with adult and fetal human kidneys, major nephron cell types were identified in D29 human kidney organoids from all four iPSC lines. Podocytes and proximal tubular cells were well developed, whereas distal tubular cells were less differentiated. At single cell resolution, we also found that the iPSCs themselves had comparable pluripotency (at D0) and modest variability at D7. At D15, organoids showed greater variability in cell proportions and a distinct pool of SOX2+ cells, suggesting that variability in mature D29 organoids is likely derived from off-target programs at a point in organoid development between D7 and D15. The pattern of variability in cell composition was then maintained through D29, driven by the off-target compartment. Importantly, organoids with higher proportions of nephron progenitor cells at D15 went on to develop more off-target cells at D29. Hence, our data reveal that future improvements in organoid protocols should focus on the time interval between D7 and D15, aiming to reduce the persistence of progenitor cells that drive the development of off-target cell populations.

Second, the applicability and faithfulness of organoids as a tool for discovery was bolstered by the fact that the vast majority of genes associated with kidney diseases was expressed in organoids across all four iPSC lines in the expected, corresponding cell types. For example, we validated the expression of MUC1 in the distal tubular compartment of D29 organoids. Mutations in MUC1 are a cause of autosomal dominant tubulointerstitial kidney disease(Kirby et al., 2013; Živná et al., 2018), a rare kidney disease without a cure. Hence, efforts to find cures for genetically defined diseases may benefit greatly from our ability to study their mechanisms in patient iPSC-derived organoids, and this study provides an important foundational step in this direction. Specifically, our work suggests that organoids derived from individual patients can indeed be used as a tool to fuel biological discovery and therapeutics. However, single cell transcriptomics using comprehensive atlases such as this may be an essential reference when trying to make meaningful comparisons between iPSC-derived kidney organoids from different patients.

Finally, we addressed the critical question of organoid quality, with a focus on reduction of off-target cells. Recent reports focused on eliminating neuronal off-target cells from kidney organoids by using an NTRK2 blocker(Wu et al., 2018). However, NTRK2 is not only expressed in off-target neuronal cells, but it is also abundantly expressed in podocytes(Caroleo et al., 2015; Li et al., 2015) in both fetal human kidneys(Metsuyanim et al., 2009) and in developing organoids (**Supplementary Fig. 18A**), raising the concern that NTRK2 blockers applied to organoid cultures may adversely affect podocyte differentiation and function. Here we identified organoid transplantation as an alternate approach to diminish off-target cells. Remarkably, organoid transplantation under the mouse kidney capsule diminished SOX2+ neuronal precursors and PMEL+ melanoma cells, in contrast to organoids grown *in vitro*, that retained a neuronal precursor SOX2+ off-target population. While some of these observations merit further studies, these data show that transplantation reduced off-target cells and improved organoid quality and maturity. Future studies will be required to test if earlier organoid transplantation, as early as D7, may eliminate off-target cells and enhance organoid formation, especially ureteric bud and collecting duct development (building on gene programs such as uniquely upregulated KLF6 and HES1 in transplanted organoids). As it stands, our protocols may benefit from incorporating organoid transplantation into rodents. Since endothelial progenitors were identified in D15 organoids, and endothelial cells were detected in D29 organoids, we can further speculate that improved vascularization may be achieved by preserving and expanding human endothelial cells in organoids in the hopes of eliminating all off-target cells and generating even more faithful, adult-kidney-like organoids.

In conclusion, these studies provide unprecedented single cell resolution into iPSC-derived kidney organoids, illuminating their reproducibility, pinpointing the source of variability, and showing that transplantation is a novel method for reducing off-target cells and improving organoid quality. This comprehensive atlas, uniquely enabled by scRNA-Seq technology, should serve as a foundational resource for the community, fueling the development of much needed therapies for patients with kidney diseases.

## Acknowledgements

We thank our colleagues in the BWH Department of Urology, Drs. Steven Chang and Dimitar Zlatev for their generous assistance with recovery of tissue from tumor nephrectomies under IRB protocol 2011P002692. We thank Kendrah Kidd and Dr. Tony Bleyer at Wake Forest Medical School for assistance in obtaining blood samples for iPSC generation. We thank the Harvard Stem Cell Institute for assistance in the generation of the iPSC cell lines N1 and N2. N1 and N2 are covered under Broad ORSP-3414 and commercially purchased lines are covered by Broad NHSR-4998. We thank our colleagues Karthik Shekhar and Matan Hofree for helpful discussions.

This work was funded by NIH grants DK095045 and DK099465 (A.G.), the Slim Initiative for Genomic Medicine in the Americas (SIGMA), and an NDSEG fellowship (E.H.S.). All RNASeq data files are deposited at GEO (Accession number GSEXXXXXX). A.G. has a financial interest in Goldfinch Biopharma, which was reviewed and is managed by Brigham and Women’s Hospital and Partners HealthCare and the Broad Institute of MIT and Harvard in accordance with their conflict of interest policies.

## Supplemental Figures

**Fig. S1.**
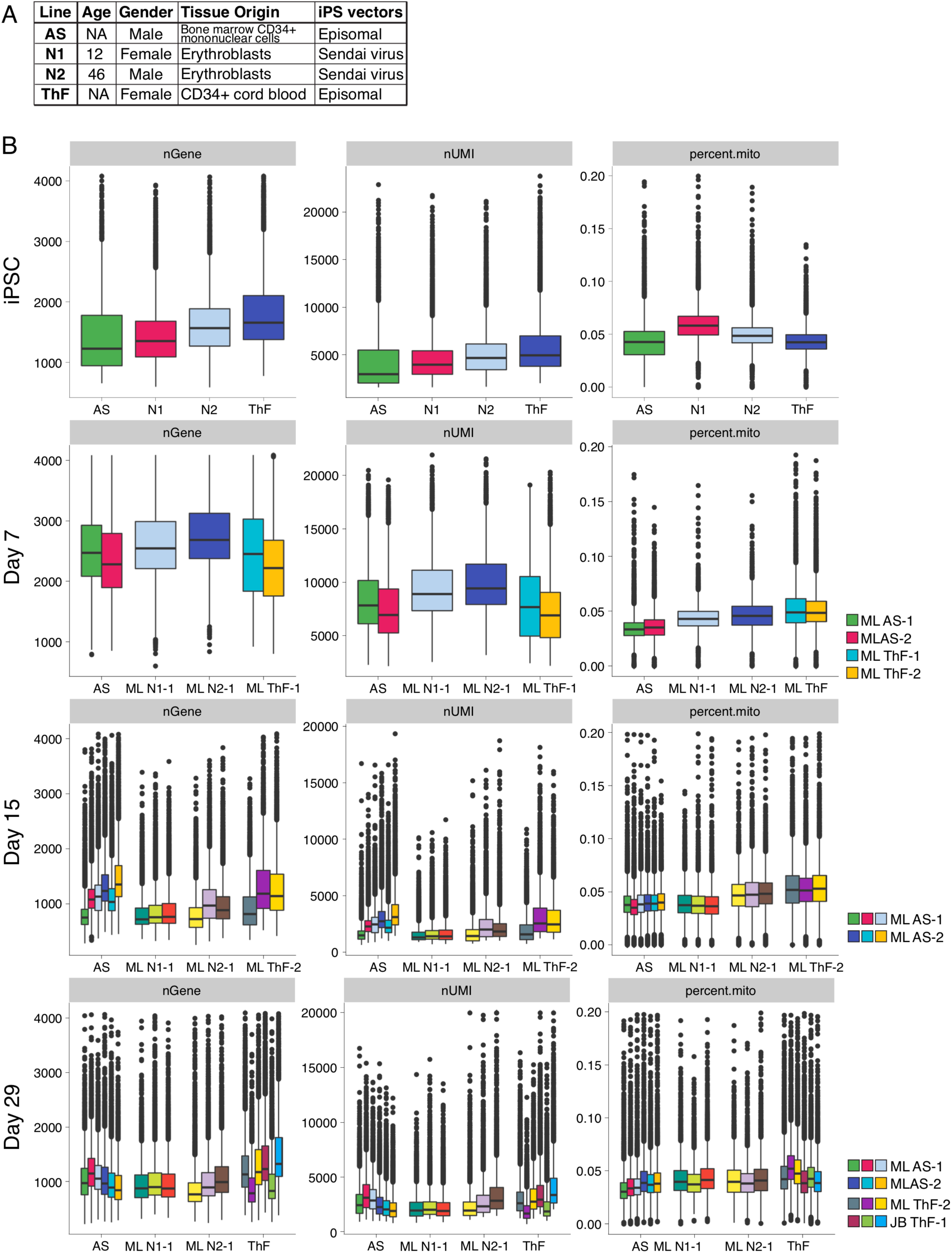
Data quality metrics for single cell analysis of kidney organoids across four iPSC lines. (A) Table of reference for 4 different iPSC lines. (B) Quality control metrics (nGene: number of genes with a normalized expression value above 0 per cell, lower QC cutoff 200; nUMI: the total number of Unique Molecule Identifiers (UMIs) detected per cell, lower QC cutoff 1000; percent.mito: The proportion of reads mapping to mitochondrial genes, upper QC cutoff of 20%) across cell lines and time points.

**Fig. S2.**
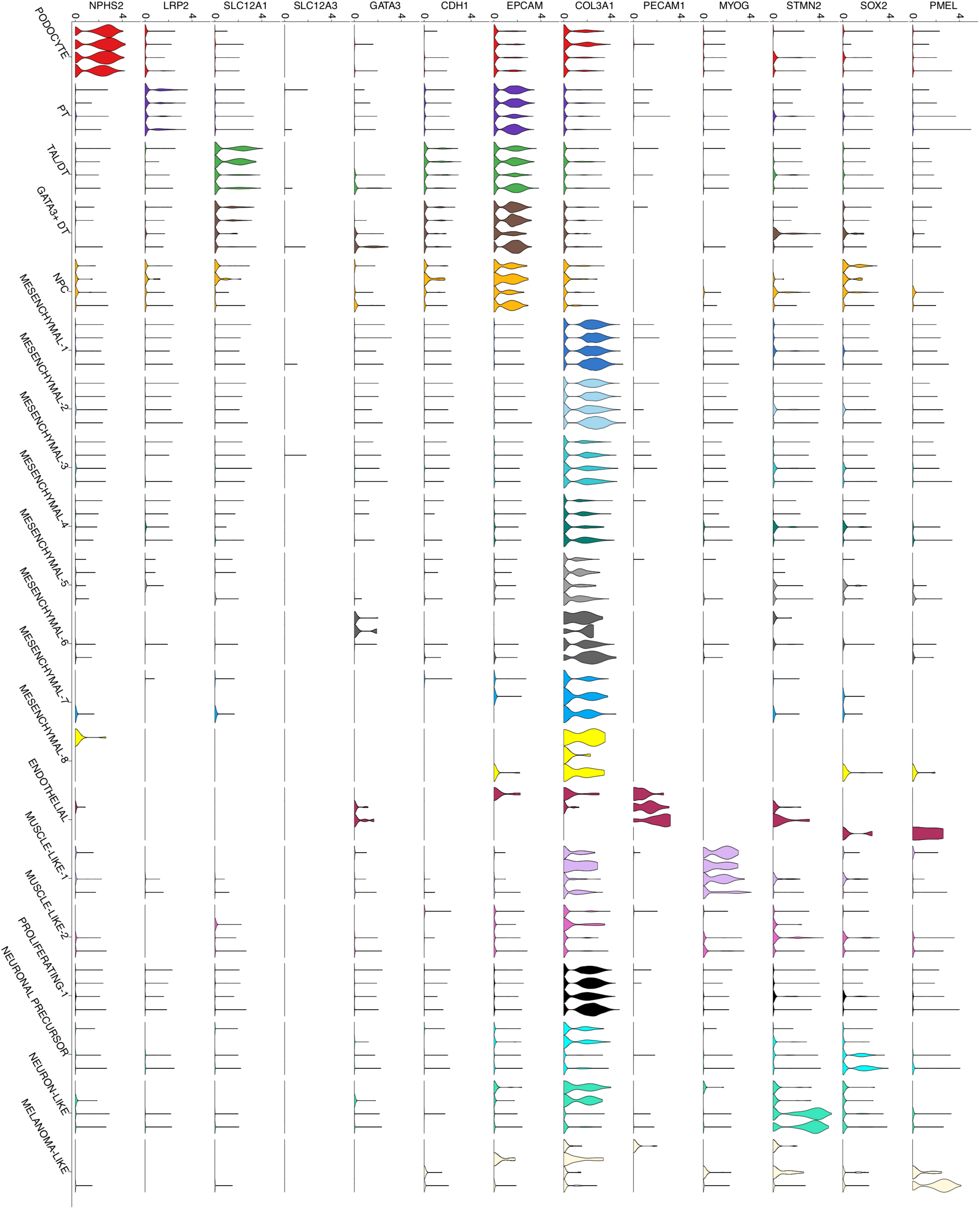
Day 29 kidney organoids express canonical markers of major nephron, mesenchymal and off-target cell types. Violin plots of single cell gene expression from day 29 kidney organoids for canonical markers. X-axis annotations represent clusters. Each violin per cluster represents a line, AS, N1, N2 and ThF in order from left to right.

**Fig. S3.**
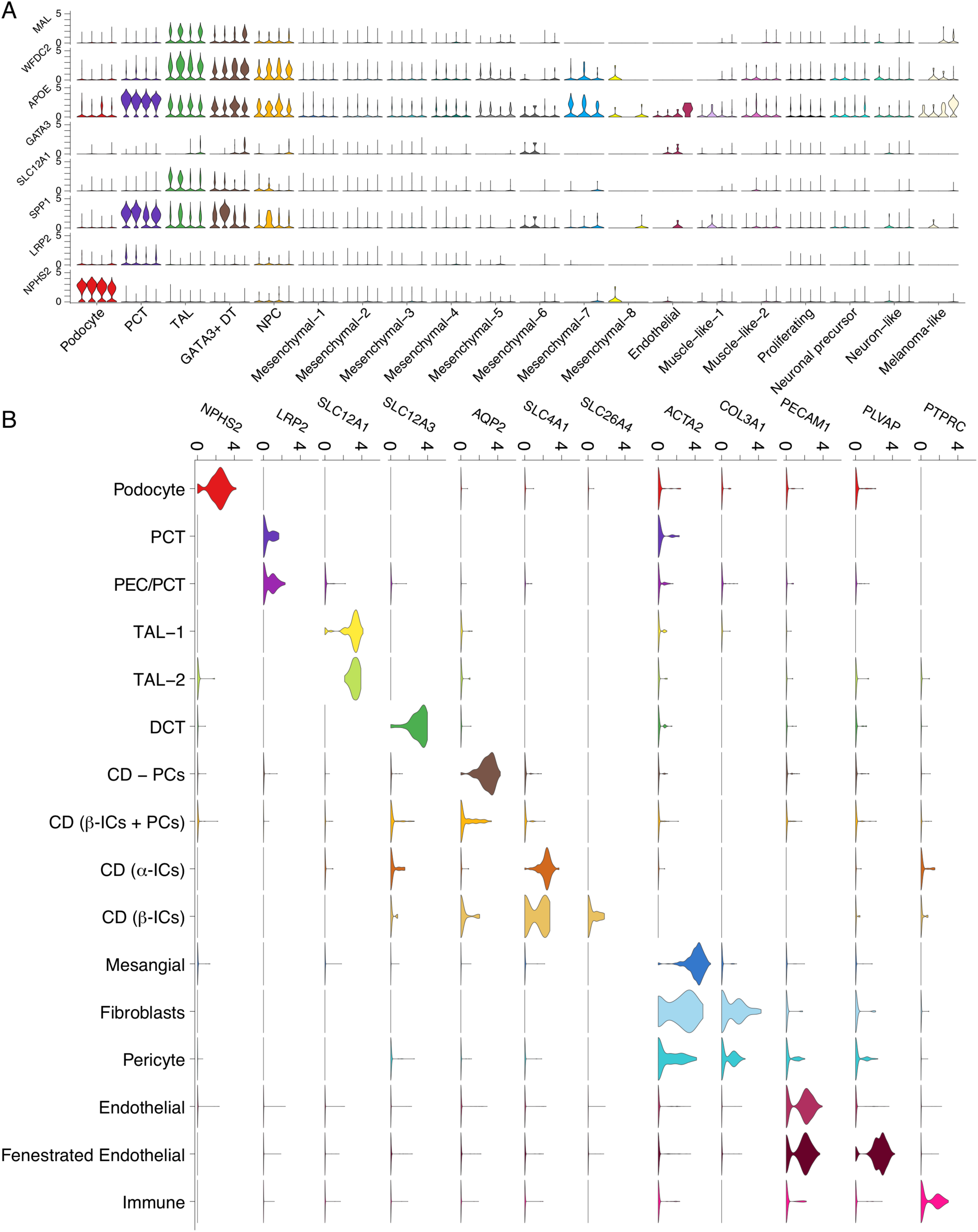
D29 data-derived markers and canonical markers of major human nephron and kidney cell types. **(A) Organoids from all lines at D29 express data-derived markers in a cluster specific manner as seen in Violin plots of single cell gene expression from day 29 kidney organoids**. (B) Violin plot of single cell gene expression from human adult nephrectomy for canonical markers. X-axis annotations represent clusters.

**Fig. S4.**
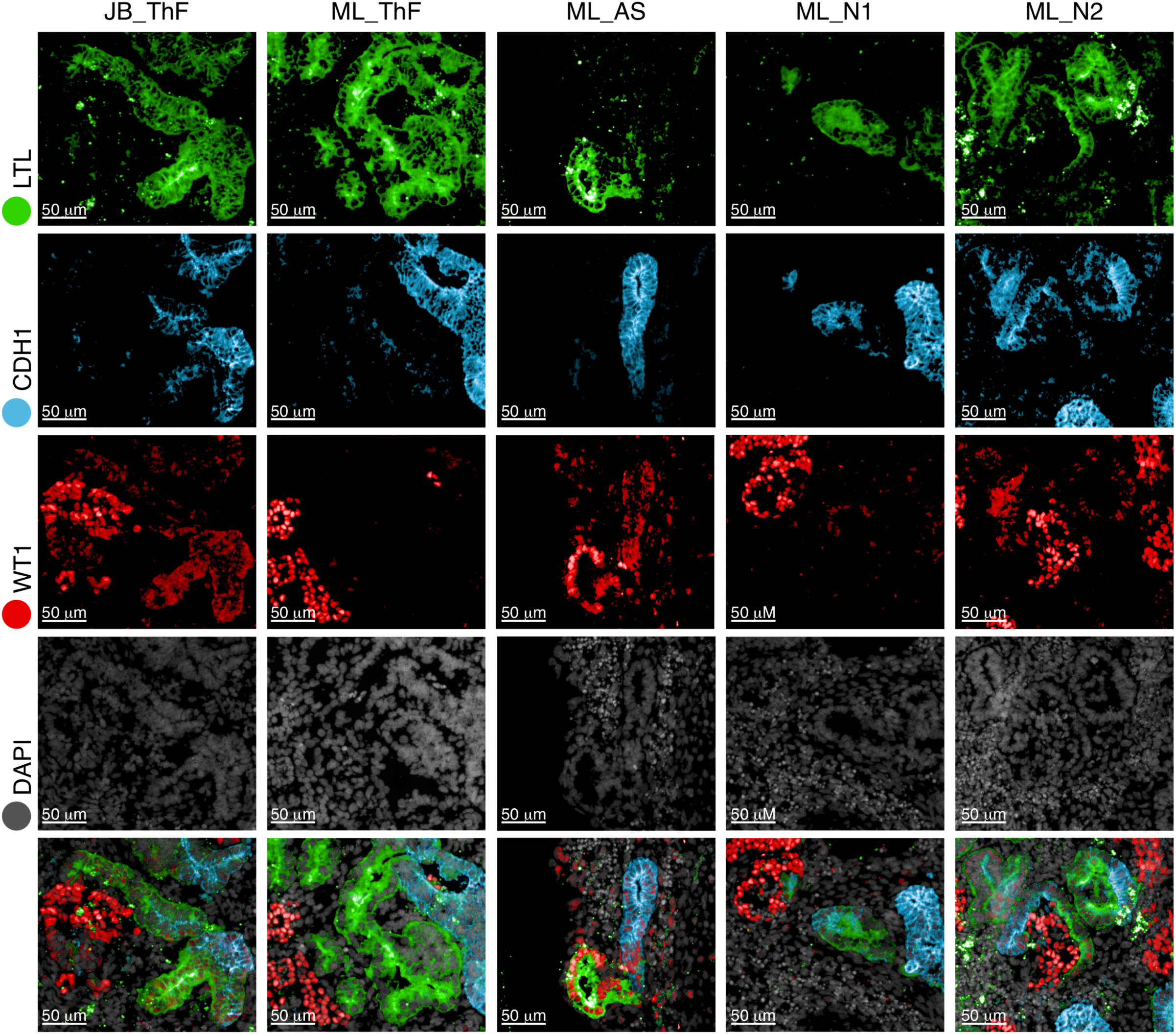
Immunofluorescence analysis of day 29 kidney organoids demonstrates presence of major nephron cell types from glomerulus to distal tubule. Immunofluorescence staining of day 29 kidney organoids for podocytes (WT1), proximal tubular cells (LTL), and the distal tubular compartment (ECAD) across two protocols (JB, ML) and four cell lines (ThF, AS, N1, N2).

**Fig. S5.**
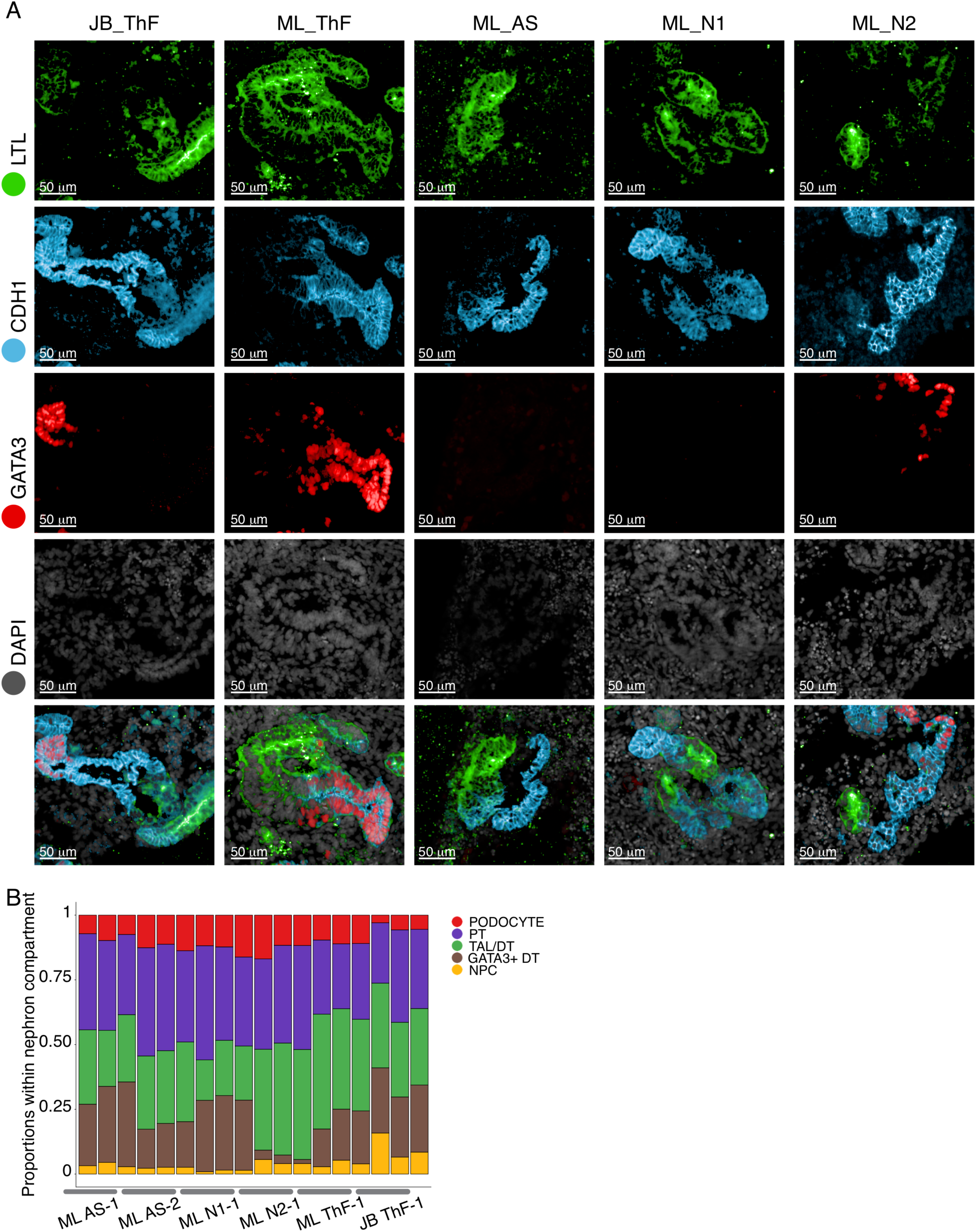
Immunofluorescence analysis of day 29 kidney organoids demonstrates presence of major kidney epithelial cell types from proximal to distal tubule. (A) Immunofluorescence staining of day 29 kidney organoids for proximal tubule (LTL), and distal nephron compartment (ECAD, GATA3) across two protocols (JB, ML) and four cell lines (ThF, AS, N1, N2). (B) Proportions of cell classes within the nephron compartment across all lines and replicates at D29.

**Fig. S6.**
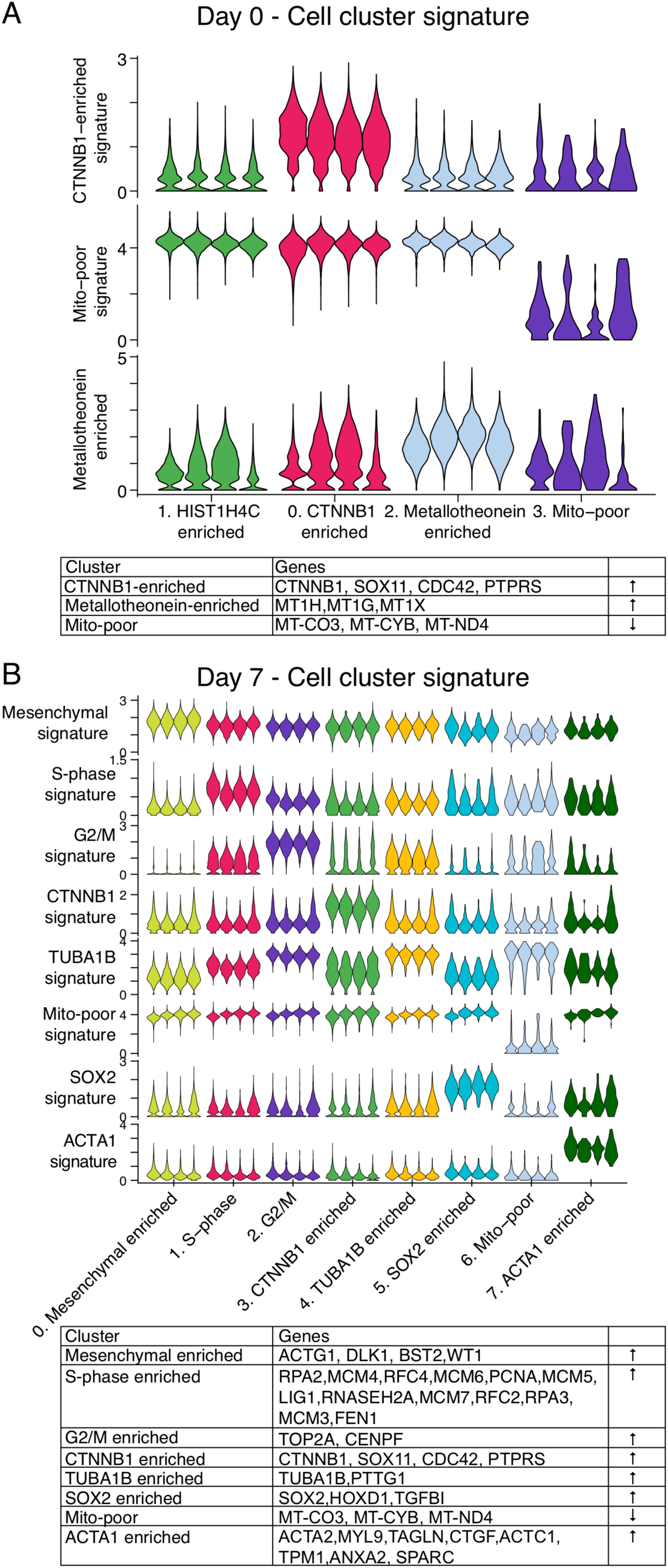
Cluster specific gene signatures for iPSC D0 and D7 stages. (A) Violin plot of average expression of cluster specific gene signatures in cells in each line at the iPSC stage. Signatures are provided in the table. (B) Violin plot of average expression of cluster specific gene signatures in cells in each line at D7. Signatures are provided in the accompanying table. In each cluster, violins for AS, N1, N2 and ThF are in order from left to right.

**Fig. S7.**
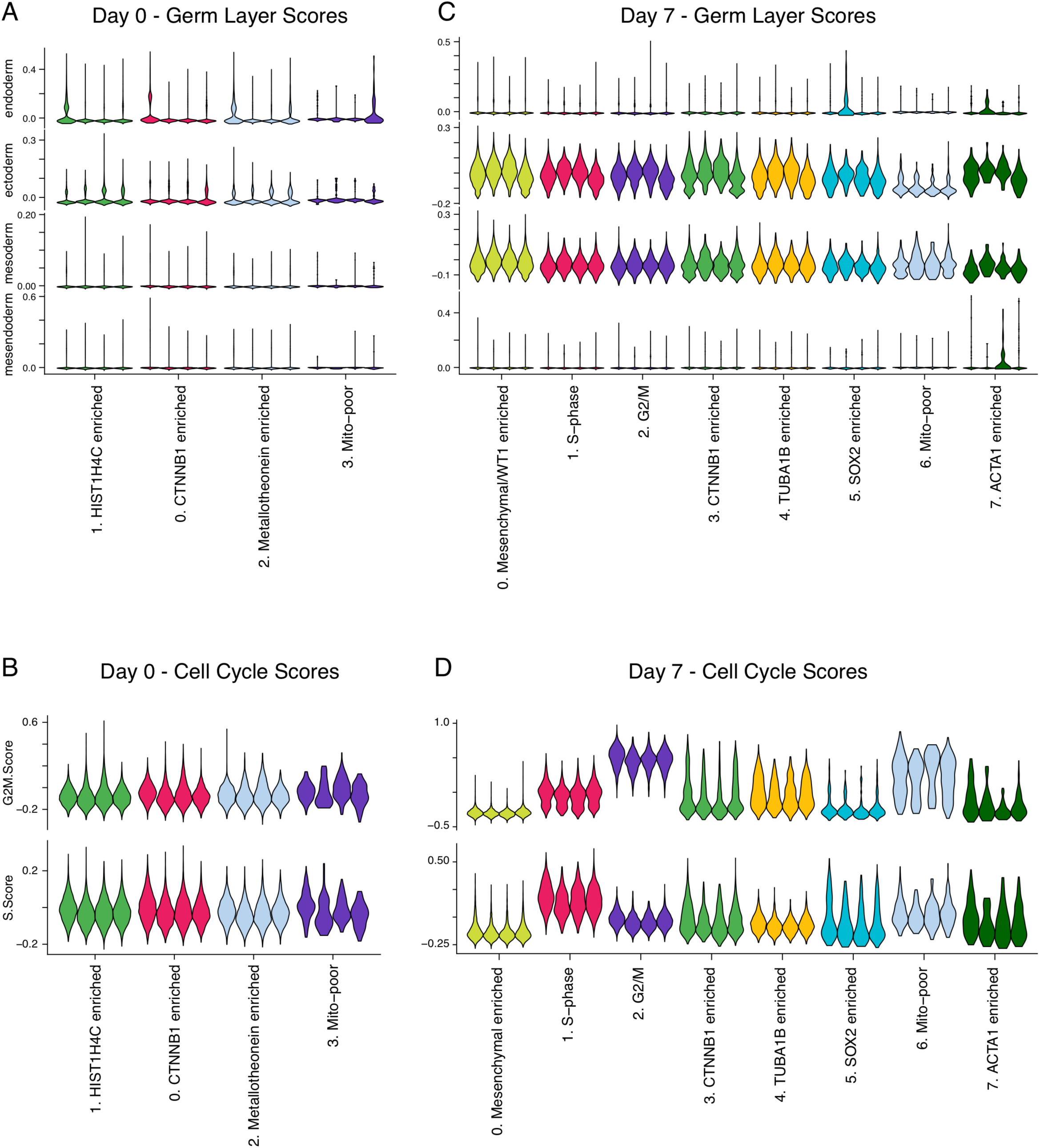
Single cell analysis of 4 iPSC lines reveals no priming for any specific germ layer and expression of cell-cycle markers. Violin plot of cluster-specific average expression of germ-layer signatures in single cells from the 4 iPSC lines at (A) D0 and (C) D7. Violin plot of scores for cell cycle phases G2M and S-phase in cells in each iPSC line at (B) D0 and (D) D7 in a cluster specific manner. In each cluster, violins for AS, N1, N2 and ThF are in order from left to right.

**Fig. S8.**
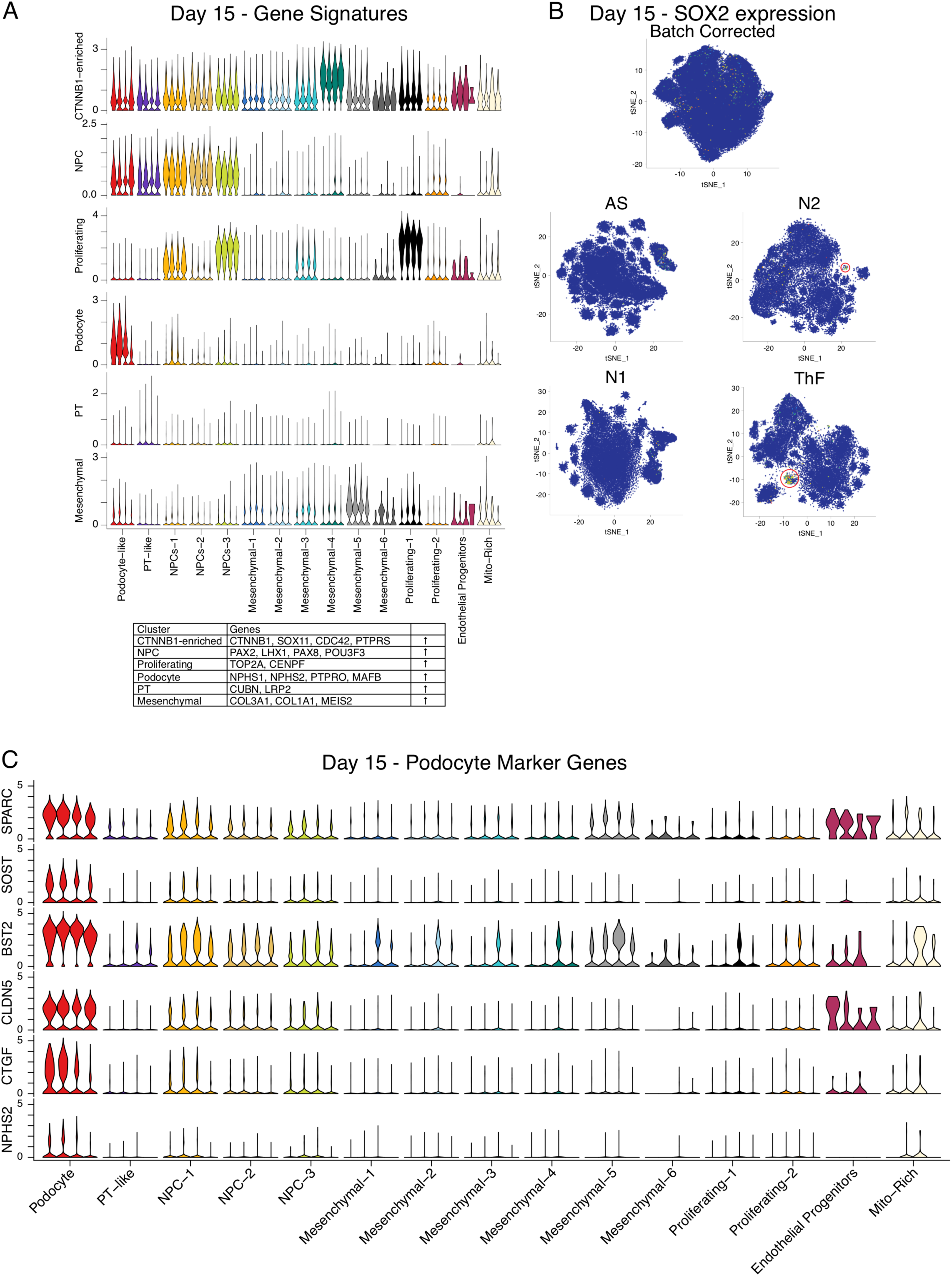
Expression of cluster specific gene signatures, line-specific SOX2+ progenitor pools and early markers of podocyte differentiation at D15. (A) Violin plot of average expression of cluster specific gene signatures in cells in each line at D15 (B) tSNE plot showing variability across the lines in presence of SOX2-positive progenitor cells at D15. Subpopulation of cells (in red circle) expressing SOX2 emerge as early as D15 in ThF and N2. (C) Violin plots from D15 organoids for expression of canonical (NPHS2) and data-driven (CLDN5, SOST, BST2, SPARC, CTGF) podocyte marker genes. In each cluster, violins for AS, N1, N2 and ThF are in order from left to right.

**Fig. S9.**
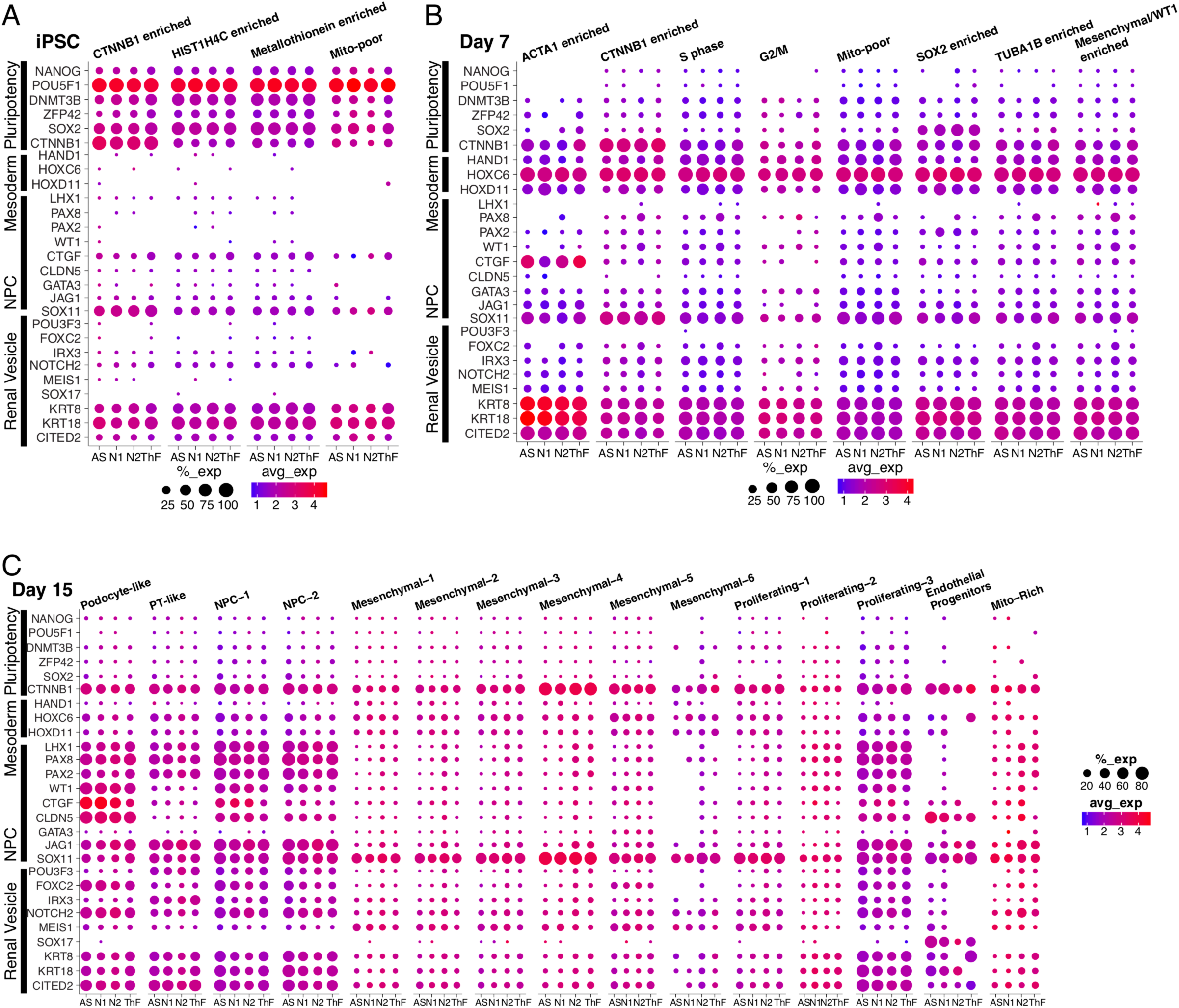
Kidney organoid differentiation follows kidney nephrogenesis as determined by expression of of transcriptional programs across organoid development time. Dot plot comparison of expression of major transcription factors and other canonical markers of nephrogenesis across organoid differentiation (iPSC, Day 7, Day 15) across the 4 iPSC lines.

**Fig. S10.**
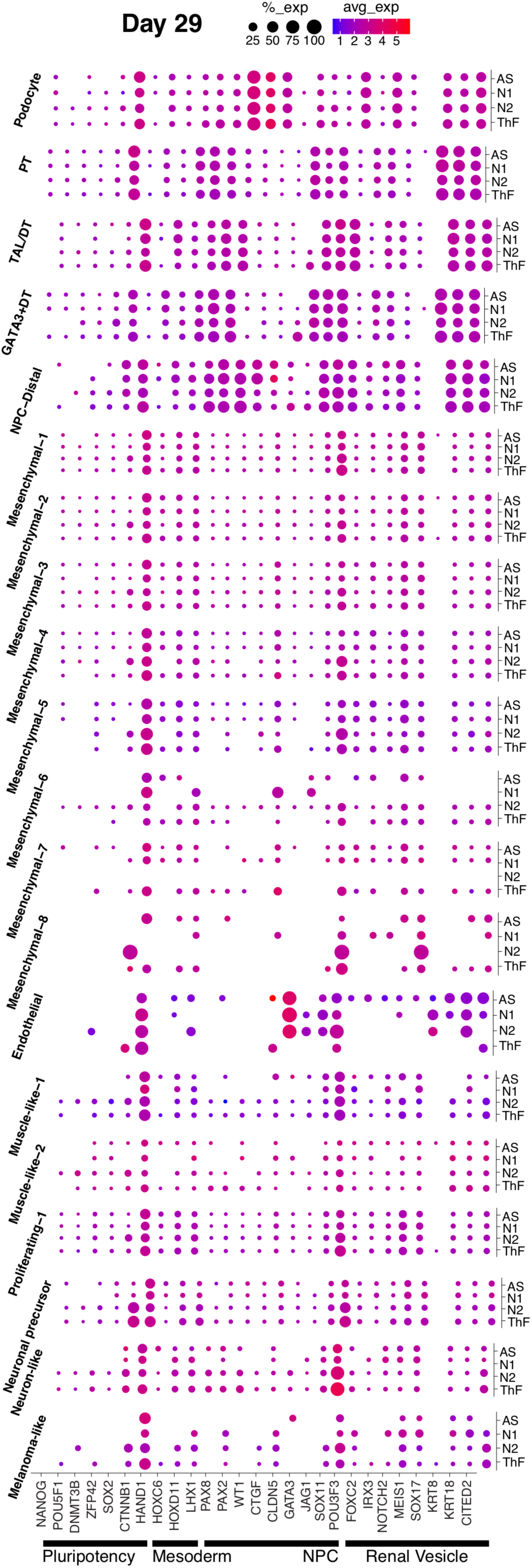
Kidney organoid differentiation follows kidney nephrogenesis as determined by expression of transcriptional programs across organoid development time. Dot plot comparison of expression of major transcription factors and other canonical markers of nephrogenesis in D29 organoids across 4 iPSC lines.

**Fig. S11.**
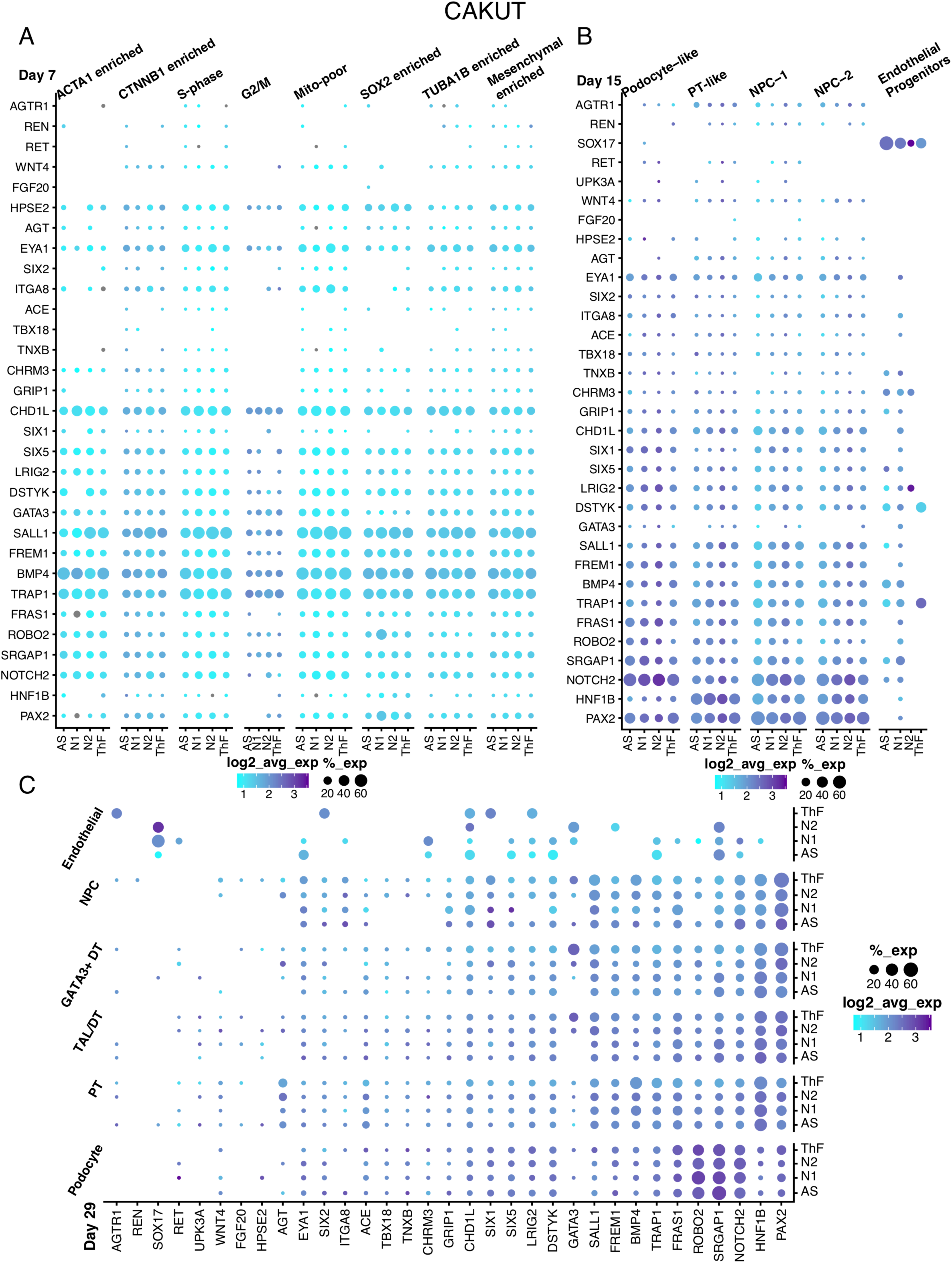
Expression of monogenic causes of congenital abnormalities of the kidney and urinary tract (CAKUT) in appropriate nephron epithelial cell types suggests utility of kidney organoids for understanding genetic kidney diseases. Dot plot comparison of gene expression in organoids across differentiation (A) Day 7, (B) Day 15, and (C) Day 29 of CAKUT-causing genes in kidney epithelial cell clusters (Day 15, Day 29) across 4 iPSC lines.

**Fig. S12.**
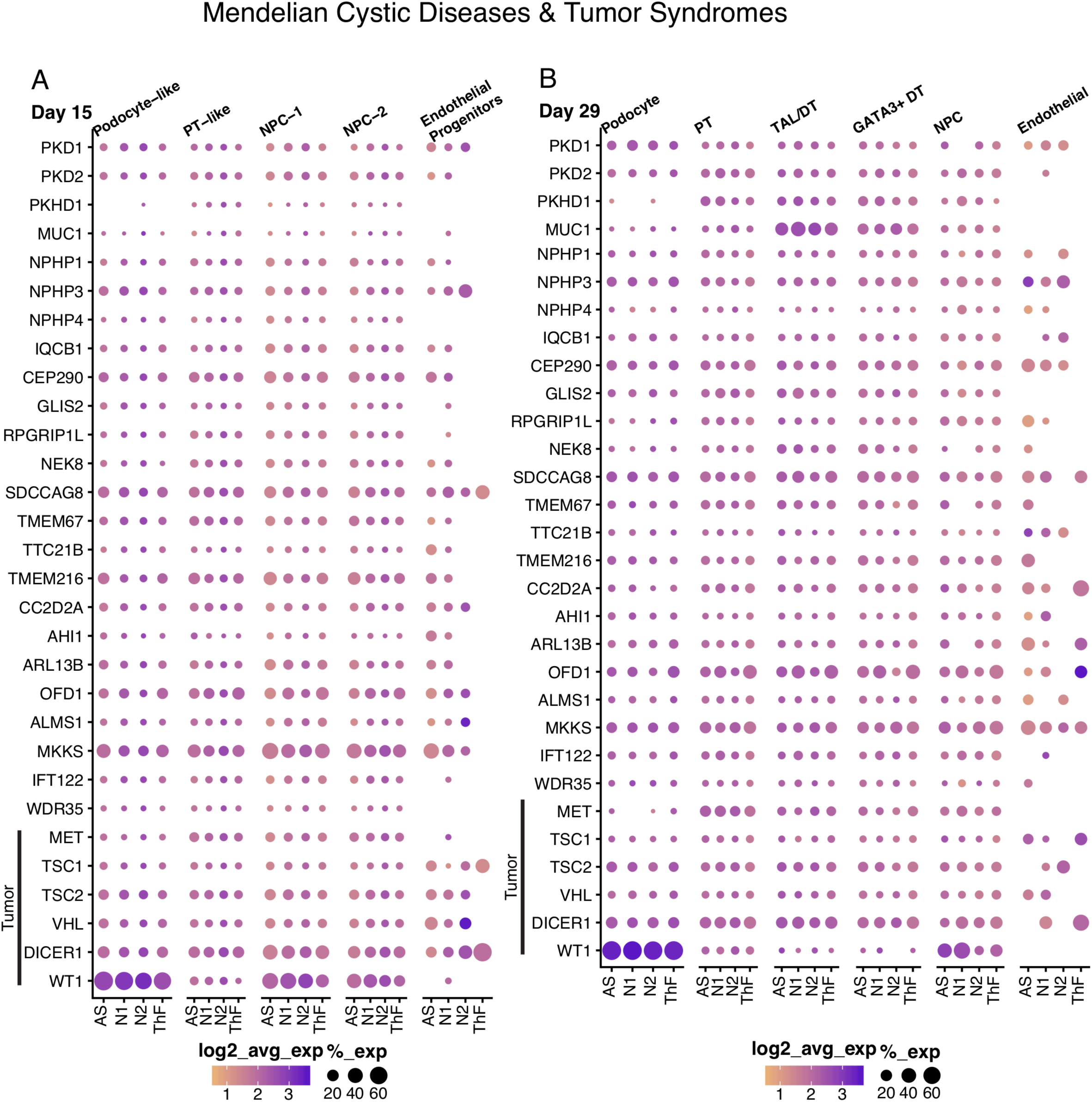
Expression of monogenic causes of hereditary renal cystic (HRC) diseases and tumor syndromes in appropriate nephron epithelial cell types suggests utility of kidney organoids for understanding genetic kidney diseases. Dot plot comparison of gene expression in organoids (A) Day 15, (B) Day 29 of HRC and tumor syndrome diseases-causing genes in kidney epithelial cell clusters across 4 iPSC lines.

**Fig. S13.**
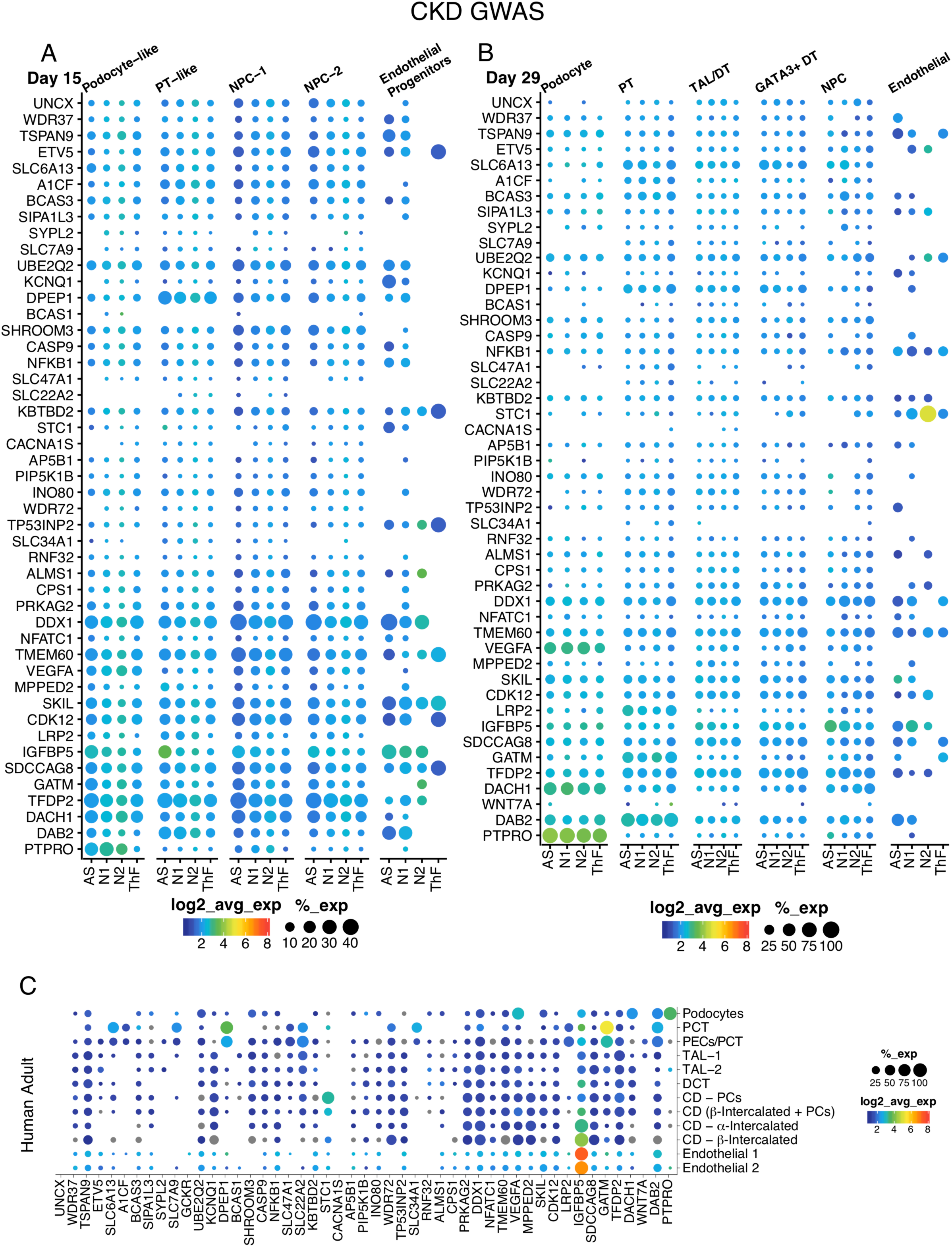
Expression of genes associated with chronic kidney diseases in appropriate nephron epithelial cell types suggests utility of kidney organoids for understanding genetic kidney diseases. Dot plot comparison of gene expression in organoids (A) Day 15, (B) Day 29) and (C) human adult of genes associated with chronic kidney disease in kidney epithelial cell clusters across 4 iPSC lines.

**Fig. S14.**
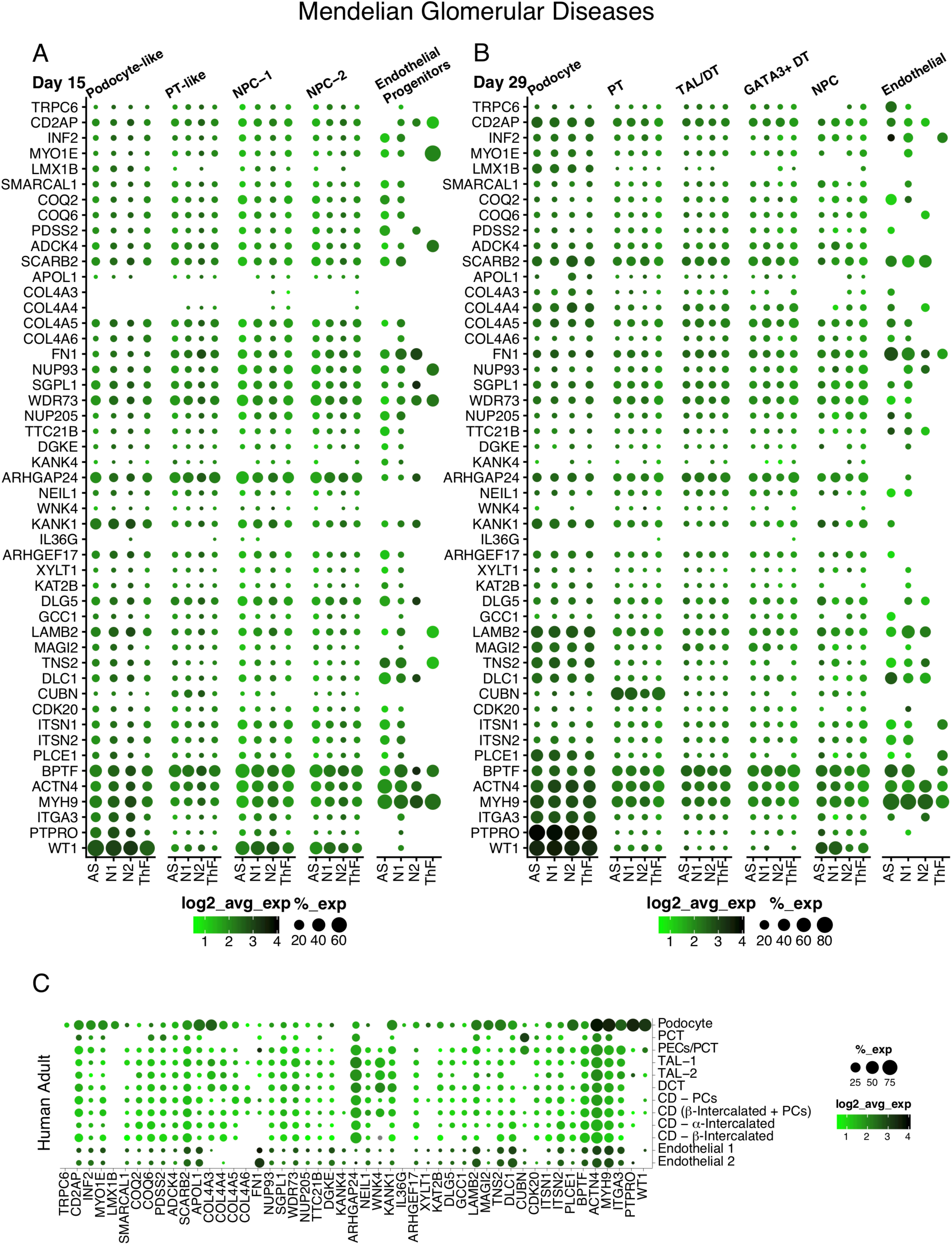
Expression of monogenic causes of hereditary glomerular diseases in appropriate nephron epithelial cell types suggests utility of kidney organoids for understanding genetic kidney diseases. Dot plot comparison of gene expression in organoids (Day 15 (A), Day 29 (B)) and (C) human adult of hereditary glomerular disease-causing genes in kidney epithelial cell clusters across 4 iPSC lines.

**Fig. S15.**
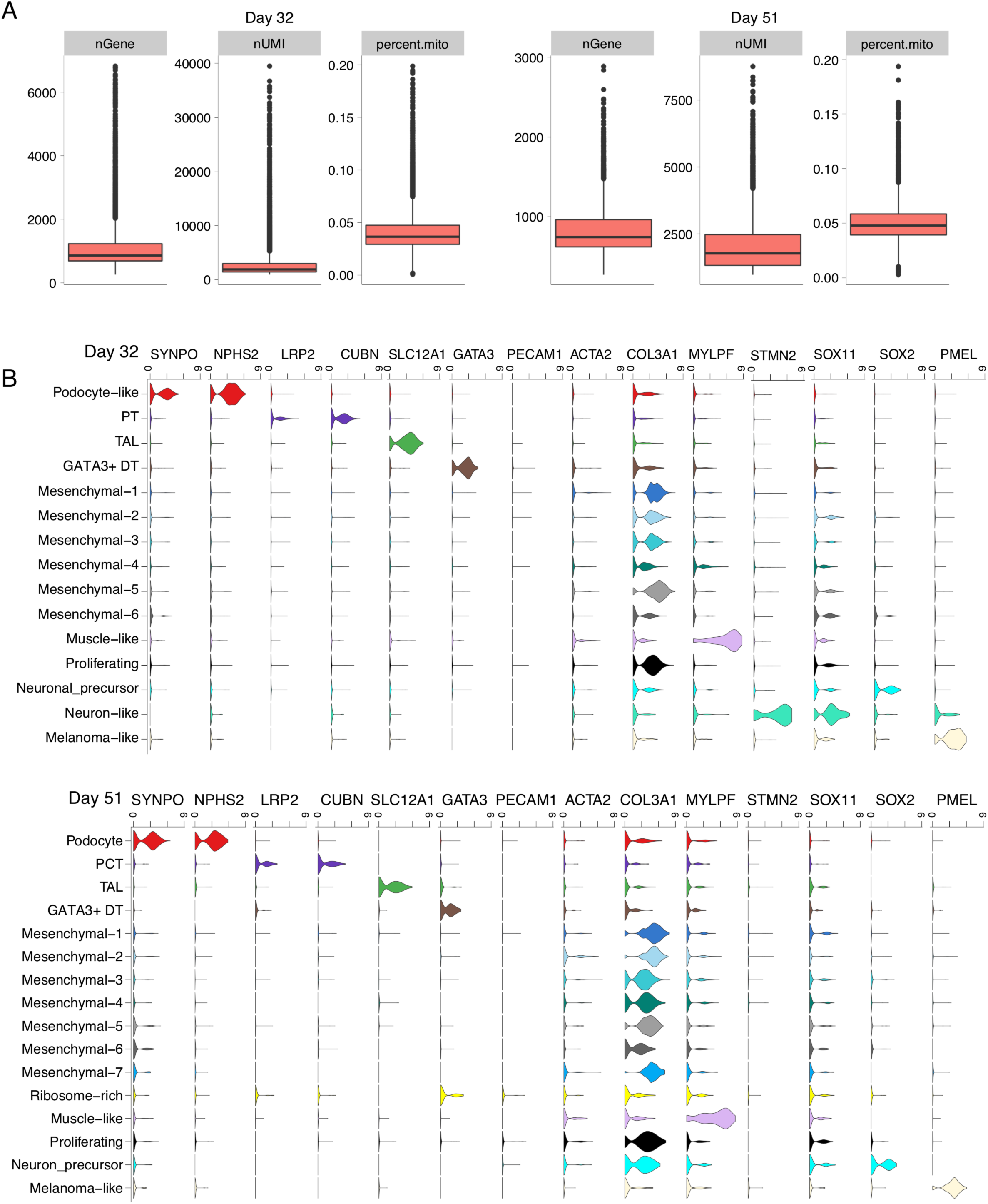
Expression of differentially enriched genes in organoids in prolonged culture found by clustering. (A) Quality control metrics of D32 and D51 control organoids. (B) Violin plot of single cells from D32 (top row) and D51 (bottom row) control organoids with gene expression of selected genes superimposed.

**Fig. S16.**
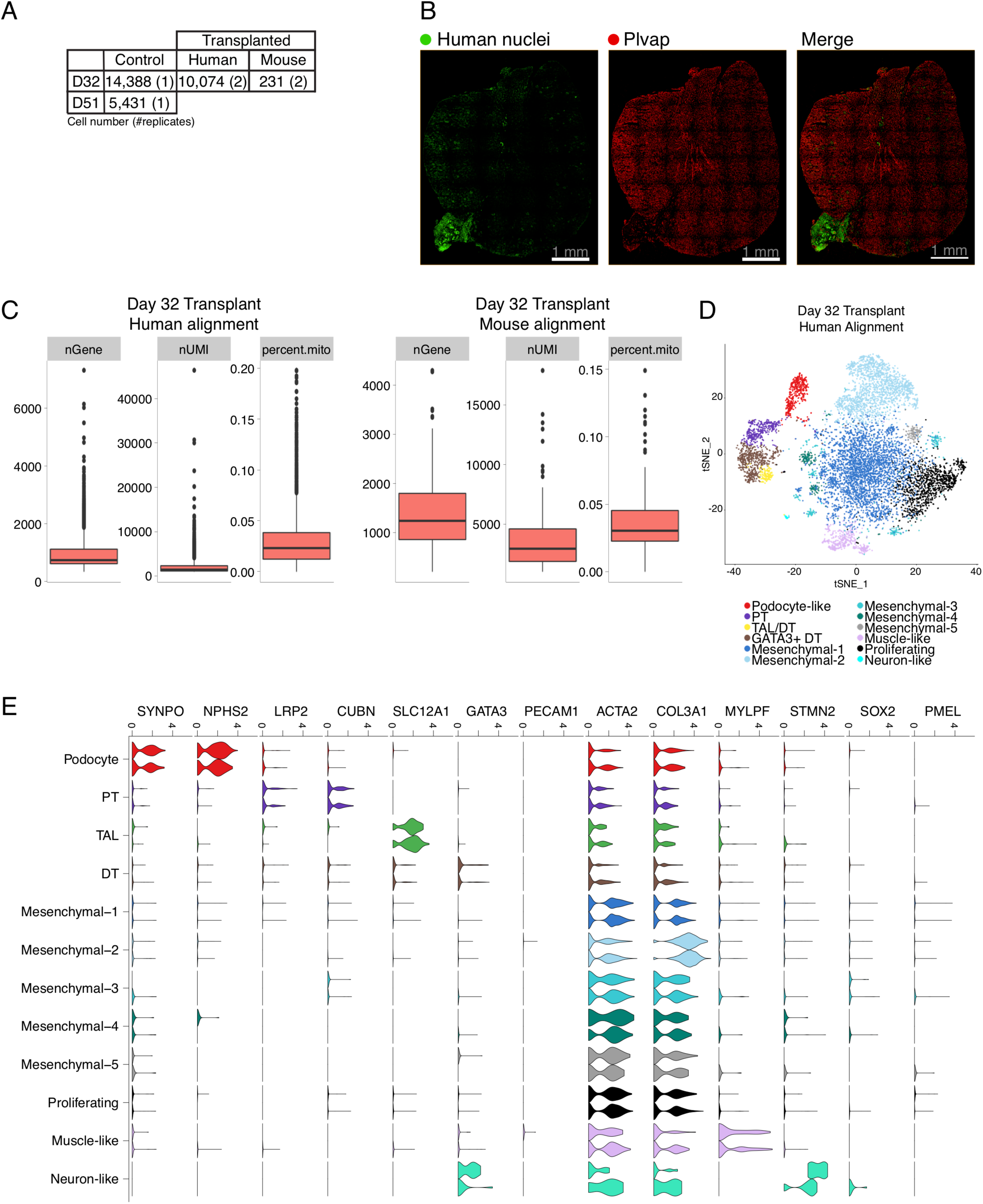
Single cell analysis of kidney subcapsular transplantation of organoids. (A) Table of cell numbers from D32 cultured and transplanted organoids. (B) Immunofluorescence staining of transplanted kidney organoids for human nuclei and endothelial cells (Plvap). (C) Quality metrics of D32 transplanted kidney organoids from alignment to the combined human and mouse transcriptomes for (left) human and (right) mouse cells. (D) t-SNE plot of human cells from D32 transplanted kidney organoids. (E) Violin plot of gene expression of canonical cluster markers in single cells from D32 transplanted organoids (human).

**Fig. S17.**
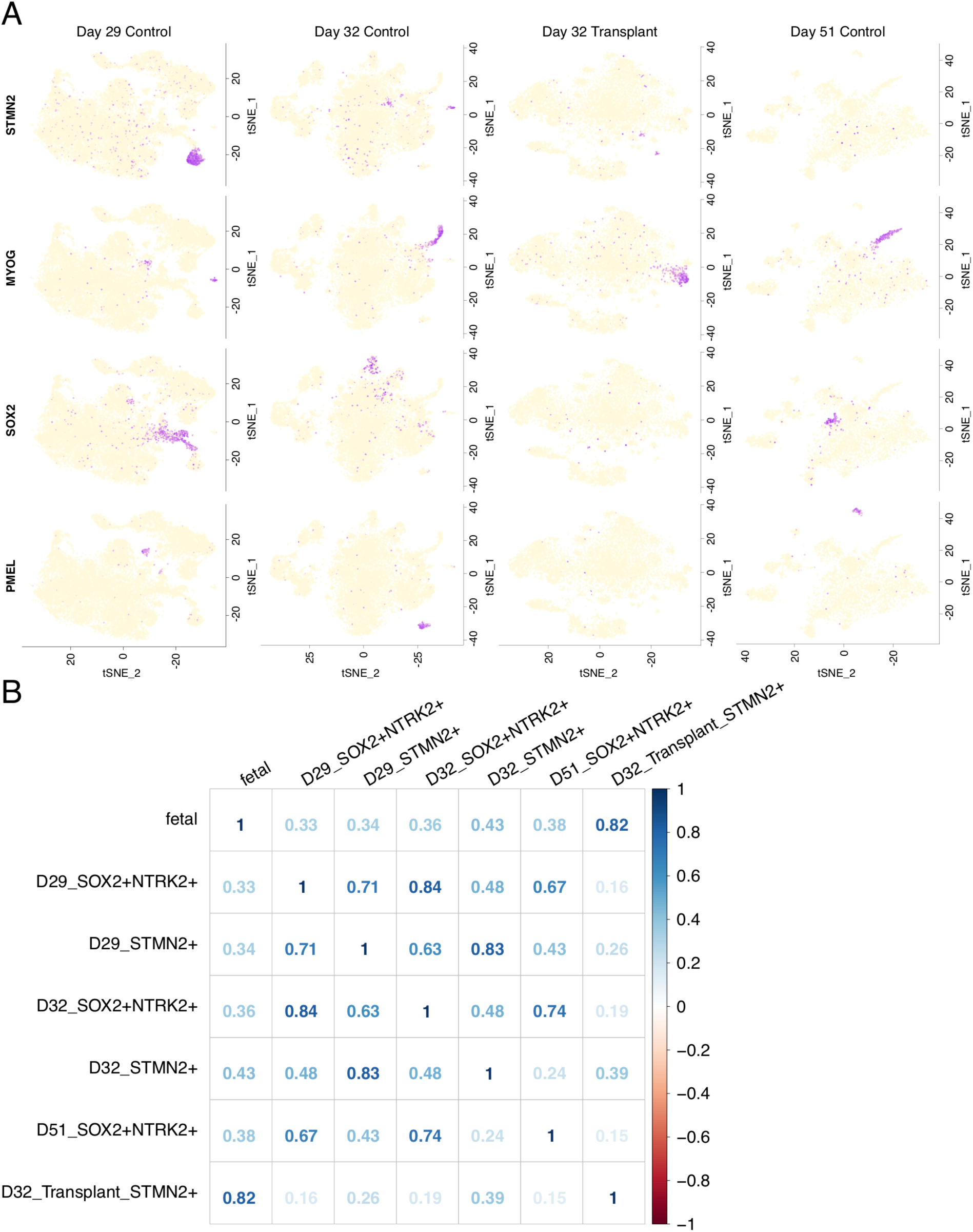
Off-target SOX2+ neuronal and PMEL+ melanoma off-target cells reduced in transplanted organoids. (A) t-SNE plot of single cells from *in vitro* (D29 and D32, and prolonged culture D51) and transplanted organoids with expression of off-target gene markers (PMEL [myeloma], SOX2 [neuronal], MYOG [muscle], STMN2 [neuronal]). The colors indicate the range of expression from low (off-white) to high (purple). (B) Spearman correlation plot indicates that STMN2+ cells in transplanted organoids correspond uniquely (ρ × 0.82) to fetal kidney, in contrast to organoids grown *in vitro*.

**Fig. S18.**
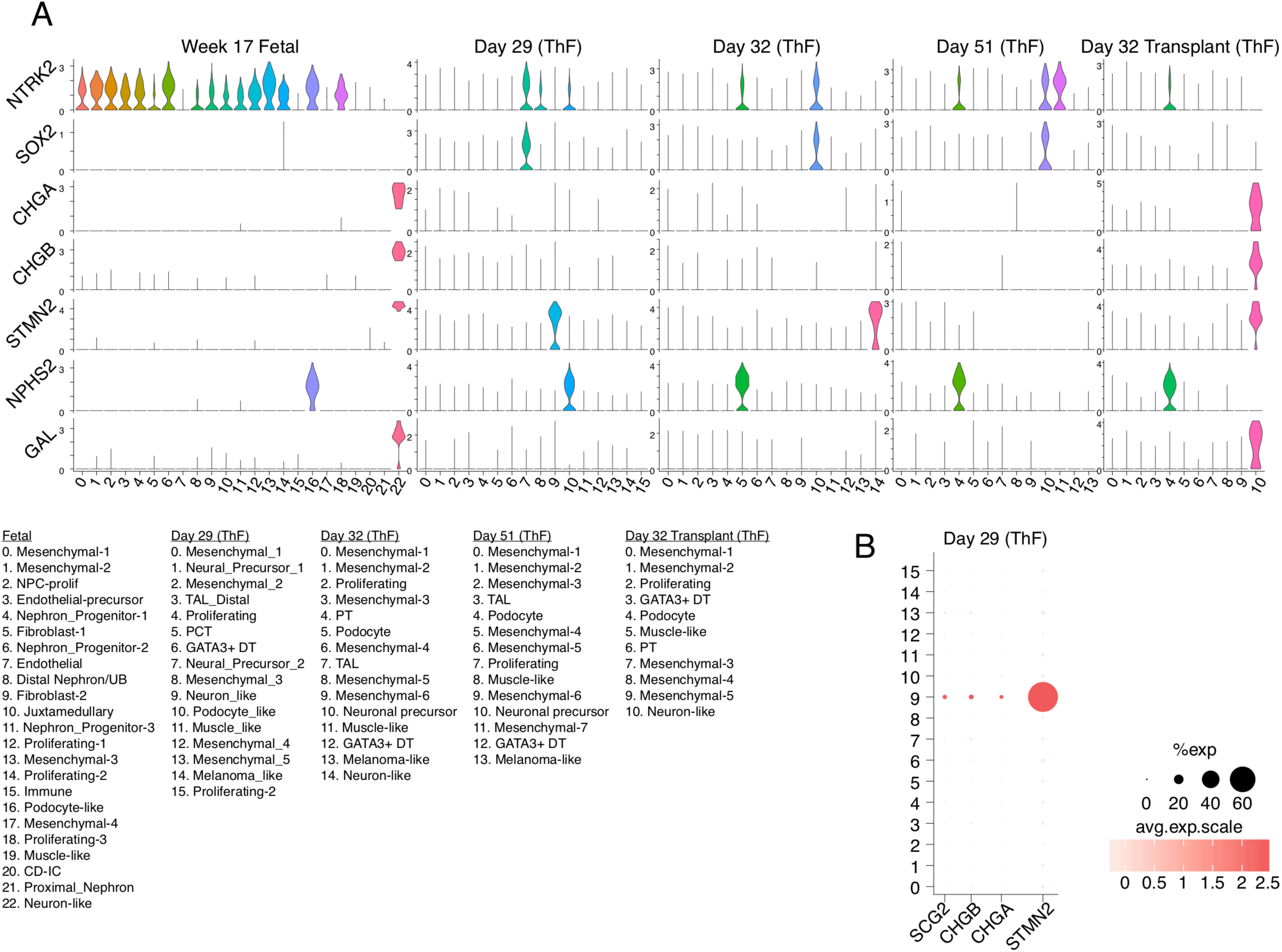
Transplanted organoids are most similar to second trimester fetal kidneys. (A) Violin plots of clusters from Day 29 and D32 controls, D51 prolonged culture and D32 transplanted showing expression of specific genes of interest. NRTK2 is abundantly co-expressed with NPHS2 (podocyte) in fetal kidney and across all organoids. The STMN2+ neuronal cluster in fetal kidney in enriched in CHGA, CHGB and GAL, and this pattern of gene expression is uniquely recapitulated in D32 transplant organoids, but not in organoids grown *in vitro*. (B) Dot plot indicates expression of CHGA and CHGB was detectable in a small number of cells in D29 STMN2+ cells.

## Materials and Methods

### Materials

*Cell Culture and Chemicals:* mTeSR1 (Stem Cell Technologies, no. 85870), Gentle Cell Dissociation Reagent (Stem Cell Technologies, no. 7174), ROCK Inhibitor Y-276*32* (Stem Cell Technologies, no. 72304), STEMdiff ™ APEL ™2 (Stem Cell Technologies, no. 05270), Accutase (Stem Cell Technologies, no. 07920), Corning Matrigel (Stem Cell Technologies, no. 354277), 6-well transwell plate (Stem Cell Technologies, no. 3450), CHIR99021 (R&D systems, no. 4423/10), Activin A (R&D systems, no. 338-AC), Human recombinant FGF-9 (Peprotech, no. 100-23), NOGGIN (Peprotech, 120-10C) and Heparin (Sigma-Aldrich, no. H4784-250mg).

*Antibodies, Immunofluorescence:* WT1 (ThermoFisher, no. PA5-16879, 1:100), ECAD, SYNPO, MUC1 (Abcam, no. ab117702, 1:500), fluorescein labeled LTL (Vector Laboratories, no. FL-1321, 1:300), GATA3, SOX2 (Cell Signaling Technology, no. 5852, 3579 1:300), Laminin (Sigma-Aldrich, no. L9393, 1:500), MEIS1 (Activemotif, no. ATM39795, 1:300) CD31 (BD Pharmingen, no. 555444, 1:300), SOX17 (R&D Systems, no. AF1924, 1:300), LRP2 (Santa Cruz Biotechnology, no. 515772, 1:100), PAX2 (Zymed laboratories, no. 71-6000, 1:300). All donkey Alexa Fluor secondary antibodies were purchased from Invitrogen (1:1000). Human nuclei (Antibodies online, no. ABIN361360, 1:300), MECA-32 (BD Biosciences, no. 553849, 1:300), NPHS1 (R&D systems, no. AF4269, 1:300) and Claudin-5 Antibody (Novus Biologicals, no. NBP2-66783, 1:300).

### iPSC culture

Human Episomal iPSC Line (ThermoFisher, no. A18945, ALSTEM, no. iPS16). N1 line (S1930 CB A) N2 line (S1973 WR I) were derived from erythroblasts using CTS™ CytoTune™-iPS 2.1 Sendai Reprogramming Kit (ThermoFisher, no. A34546) at Harvard Stem Cell Institute (HSCI) iPS Core Facility. The N1 and N2 cell lines were characterized for pluripotency and spontaneous differentiation to the three germ layers using qPCR based on standard protocols at the HSCI Core Facility. All iPSC cultures were maintained in mTeSR1 medium in T25 flasks coated with Matrigel. Cells were passaged using Gentle Cell Dissociation Reagent. All lines were confirmed to be karyotype normal and maintained below passage 15 and all the cell lines were routinely checked and were negative for mycoplasma.

### Kidney Organoid Differentiation

Kidney organoids were generated following the protocol described by Takasato et al., 2016 (Takasato et al., 2016) or Morizane et al., 2017 (Morizane and Bonventre, 2017) with slight modifications:

Takasato et al., 2016: 375K iPS cells were plated in a T25 flask in mTeSR1 media and ROCK Inhibitor, Y-27632 (10 uM). After 24 hours, cells were treated with CHIR99021 (8 uM) in APEL2 medium for four days, followed by recombinant human FGF-9 (200 ng/mL) and heparin (1 ug/mL) for an additional three days. At day seven, cells were dissociated into single cells using ACCUTASE™. 500K cells were pelleted at 350xg for 2 min (twice with 180° flip) and transferred onto a 6-well transwell membrane. Pellets were incubated with CHIR99021 (5 uM) in APEL2 medium for one hour at 37°C. After, the medium was changed to APEL2 supplemented with FGF-9 (200 ng/mL) and heparin (1 ug/mL) for an additional five days, and an additional two days with heparin (1 ug/mL). The organoids were maintained in APEL2 medium with no additional factors until day 29-51 for downstream experiments. Medium was changed every other day.

Morizane et al., 2017: iPS cells were plated as described above. After 24 hours, cells were treated with CHIR99021 (10 uM) and NOGGIN (5 ng/mL) in APEL2 medium for four days, followed by Activin A (10 ng/mL) for two days, and FGF-9 (10 ng/mL) for an additional two days. At day 8, the cells were dissociated and transferred to a 6-well transwell plate as described above. Pellets were incubated with CHIR99021 (3 uM) and FGF-9 (10 ng/mL) in APEL2 for two days, followed by four days with only FGF-9 (10 ng/mL). Organoids were maintained in APEL2 medium with no additional factors until harvest at day 2*5*-28 for downstream experiments. Medium was changed every other day.

### Single cell isolation for 10X genomics

Day 28 mature kidney organoids were washed twice with PBS and incubated with Accumax (Stem Cell Technologies, no. 07921) for 10 minutes at 37°C and were dissociated into single cells using a 27G syringe (BD biosciences, no. 305540). Cells were spun down at 350xg for 5 minutes, resuspended in PBS, filtered through a 40µm filter (Corning, no. 352340), and checked for viability.

### Library preparation and single cell sequencing

Single cells were processed through the 10X Chromium 3’ Single Cell Platform using the Chromium Single Cell 3’ Library, Gel Bead and Chip Kits (10X Genomics, Pleasanton, CA), following the manufacturer’s protocol. Briefly, 10,000 cells were added to each channel of a chip to be partitioned into Gel Beads in Emulsion (GEMs) in the Chromium instrument, followed by cell lysis and barcoded reverse transcription of RNA in the droplets. Breaking of the emulsion was followed by amplification, fragmentation and addition of adapter and sample index. Libraries were pooled together and sequenced on Illumina HiSeq.

### Immunofluorescence

Organoids were fixed in 4% paraformaldehyde (Alfa Aesar, no. J61899-AP), cryoprotected in 30% sucrose solution overnight, embedded in optimum cutting temperature (OCT) compound (VWR, no. 25608-930), flash frozen in dry ice with ethanol, and kept at −80°C overnight. Organoids were cryosectioned (Leica CM1950 Clinical Cryostat) at 6□m and mounted on Micro Slides, Superfrost™ Plus (VWR, 48311-703). Slides were washed with PBS (1 time, 5 minutes), blocked for 20 minutes (5% normal donkey serum, 1.5% Tween-20), and incubated overnight at 4°C with primary antibody (in blocking buffer). After, the slides were washed with PBS (3 times, 10 minutes each) and incubated with secondary antibody (PBS, 1.5% Tween-20) for two hours at room temperature. The slides were then washed with PBS (1 time, 10 minutes). For phalloidin staining, the cells were incubated with Phalloidin-647 (ThermoFisher, no. A22287, 10U/mL in 1.5% Tween-20) for 20 minutes at room temperature and washed with PBS (1 time, 10 minutes). The sections were stained with DAPI (ThermoFisher, no. 62248, 1: 10,000 in PBS) for 5 minutes, washed with PBS (3 times, 10 minutes), and mounted using ProLong™ Gold antifade reagent (ThermoFisher, no. P36930). Images were obtained by confocal microscopy (PerkinElmer Opera Phenix High Content Screening System).

### Animal Experiments

Animal experiments were done at Custom contract research company Biomere (Biomedical Research Model company (https://biomere.com). Biomere has all the IACUC approval for animal experiments. Recipient mice (n × 8, NOD scid gamma (NSG) 8-week-old female mice, The Jackson Laboratory, no. Jax # 005557). Mice were anesthetized with ketamine/dexdomitor and for pain relief animals were injected with burenorphine. ThF kidney organoids generated using ML protocol (Day 7+11 and Day 7+18) were transplanted onto the left kidney capsule of mice as described in Cathelijne et al., 2018 (van den Berg et al., 2018). Day 7+11 organoids were collected after 14 days of implantation and Day 7+18 organoids were collected after 26 days of implantation for single cell RNA sequencing, Immunohistochemistry and for Electron Microscope (EM) analysis.

### Single cell isolation from human kidney tissue

Samples of macroscopically normal cortex were obtained from a tumor nephrectomy specimen, distant from the tumor site and after appropriate patient consent, in accordance with IRB and institutional guidelines. Tissue was cut into 1mm x 1mm cubes and placed in 0.25mg/ml liberase TH (Roche Diagnostics, Indianapolis, USA) dissociation medium. Following further dissection, the tissue was incubated at 37C for 1 hour in a thermomixer at 600rpm. Samples were regularly triturated during the incubation period using a 1ml pipette, after which 10% heat-inactivated FBS RPMI was added to stop the digestion. Centrifugation at 500g for 5 minutes at room temperature and removal of the supernatant was followed by the addition of ACK lysing buffer to remove erythrocytes (Thermo Fisher Scientific, Waltham, USA). Repeat addition of ACK lysing buffer was performed given the kidney was not perfused prior to removal. Following centrifugation, the resulting cell pellet was incubated with Accumax at 37C for 3 minutes (Innovative Cell Technologies Inc, San Diego, USA). 10% FBS RPMI was again used to neutralize the accumax and centrifugation was followed by resuspension of the cell pellet with 0.4% BSA/PBS. The single cell suspension was then filtered using a 30um CellTrics filter (Sysmex America Inc, Lincolnshire, USA) with the resulting cell concentration and viability determined using trypan blue. 10,000 cells were then loaded into the 10x Genomics microfluidic system according to the manufacturer’s guidelines (10x Genomics, Pleasanton, USA).

## Computational Methods for Data Analysis

### Study Design

Replicates from both sexes were incorporated wherever possible. Organoids were pooled on lanes using a randomized design to ensure that organoids replicates from an individual batch (donor, replicate, condition) were distributed across lanes.

### Preprocessing of 10x droplet based sequencing outputs

We used the *Cellranger* toolkit (v2.1.1) to perform de-multiplexing using the “cellranger mkfastq” command, and the “cellranger count” command for alignment to the human transcriptome, cell barcode partitioning, collapsing unique-molecular identifier (UMI) to transcripts, and gene-level quantification.

Quality Control: We filtered cells to only include barcodes with minimum mapped UMIs of 1000, summarizing to at least 200 genes for downstream analysis. Further, the percentage of reads mapping to mitochondrial genes was capped at 20%.

### Inferring cell-types from individual donor organoid, fetal and adult kidney cells

We used the default settings in the *Seurat* R package(Butler et al., 2018) (v2.3) for normalization (NormalizeData) of the gene expression counts and identifying variable genes (FindVariableGenes). Briefly for normalization, UMI counts in each gene in a cell are divided by the total UMI count for the cell, divided by a factor of 100000 and log-transformed to obtain log(TPX+1) values. After mean-centering, we performed dimensionality reduction using Principal Component Analyses (RunPCA) on the highly variable genes computed in the previous step. We retained 50 PCs for unbiased clustering (FindClusters) by building a k-nearest neighbor graph and smart-moving local community detection algorithm. For iPSCs, the first 8 Principal Components sufficiently captured all the variance. The resolution parameter was adjusted as needed: for the human adult sample, resolution of 3 recovered expected renal cell compartments, for day 28 and 15, resolution of 1 was used. We computed an embedding of the data in 2-D space using t-distributed stochastic neighbor embedding (tSNE) in the PC space for visualization (RunTSNE), independent of the clustering step.

### Assignment of cell-identity and gene-set signatures

Cluster-enriched or marker genes were computed using the Wilcoxon-Rank sum test (FindAllMarkers) for differential expression of genes in the cluster cells vs all other cells, and selecting those genes that pass the adjusted p-value (Benjamini-Hochberg(Benjamini et al., 1995) FDR) cutoff of 0.05 as cluster-representative. Cluster identity was assigned by comparing data-driven genes with a list of literature-curated genes for kidney developmental and mature cell types. Sub clustering was performed when a single cluster represented marker genes from multiple renal epithelial cell-types. We checked that cluster membership was not exclusive to a single replicate. Cells were scored for embryonic germ layer-specific gene signatures using Seurat’s “AddModuleScore” function. Seurat’s “CellCycleScoring” function was used to score cells for different stages of cell-cycles.

### Curation of gene signatures

The cell-cycle signatures were obtained from Tirosh et al. The germ-layer signatures were obtained from Tsankov et al. 2015. Only those genes were included which passed the cutoff of. We curated a set of genes, both transcription factor or not, critical for early nephrogenesis from the kidney development literature.

### Combined analyses of organoids from multiple donors

Clusters, particularly representing interstitial cell types, segregated by donor origin when default analyses as described above was performed. To identify similar cell types among organoids from multiple donors, we co-embedded the cells using canonical correlation analyses using Seurat’s “MultiCCA” function. 20 canonical components were used for clustering. Clusters were assigned identity based on expression of literature-curated marker genes as well as verified by differential expression analysis for individual lines (AS or ThF).

### Analysis of Mouse transplants

For cells from the mouse transplants, we aligned the sequenced reads to a reference combining the human and mouse transcriptomes using the Cellranger software as described before. Multimapped reads were discarded and two expression matrices, representing mouse and human barcodes were derived. In case of barcodes that remained in both matrices after quality filtering, we assigned their identities based on the transcriptome that yielded the higher number of total UMIs. Downstream analysis was performed as described above.

### Analysis of human fetal data

Trimester 1 human fetal kidney single-cell transcriptomes were downloaded from the Data Supplement in Trimester 2 Human fetal kidney single-cell transcriptomes were downloaded from NCBI GEO GSE112570(Young et al., 2018). In both cases, the data were available in the format of gene expression count matrices which were directly plugged into cell-type identification and assignment routines as described above.

### Assessment of compositional differences

The Jenson-Shannon Divergence was computed between any two vectors of cell-proportions using the “JSD” function in the *philentropy* R package, using the default log2 transformation. The AS, passage 1 (D7, D15, D29) was excluded when computing summary statistics as it was an outlier in terms of extremely low nephron numbers. ThF, passage 1 (D7) was also excluded as it did not undergo nephrogenesis.

### Comparison of cell-types between organoids and human kidney cells

We used the R package randomForest (Liaw, A and Wiener, 2002) to train a multi-class Random Forest classifier (5000 trees) on the mature ThF organoids. The number of training cells per cell-type was set at 1000 or 70% of cluster membership, whichever was lower. The remaining cells comprised the test set. The classifier was then used to predict cell types in fetal kidney cells derived from individual kidneys at trimesters one and two, and adult human kidney cells. We inferred cell-types for fetal and adult cells using unbiased clustering as described earlier. Prediction outcomes and concordance of cell-types were visualized by a dotplot representation, where x-axis levels were ThF cell-types and y-axis levels were input (fetal or adult) cell-types. Each dot (size and color) represented the percentage of cells in the input (y-axis level) cell-types predicted to be a ThF (x-axis level) cell-types. The supervised classification analyses revealed high concordance between organoid and human kidney nephron epithelial and podocyte cell types.

### Plotting and visualization

The R package *ggplot2*(Wickham, 2016) was used for visualization of tSNEs, cell-type proportions, boxplots and dotplots. In the dotplot representation, each dot size represented the percentage of cells and the color represented the average nonzero gene expression in log2 scale. For Fig 4A-C, we only display dots that represent expression in at least 5% of the respective cluster.

## References

Araki, S.I. (2014). APOE polymorphism and diabetic nephropathy. Clin. Exp. Nephrol.

Ashraf, S., Kudo, H., Rao, J., Kikuchi, A., Widmeier, E., Lawson, J.A., Tan, W., Hermle, T., Warejko, J.K., Shril, S., et al. (2018). Mutations in six nephrosis genes delineate a pathogenic pathway amenable to treatment. Nat. Commun. 9.

Bantounas, I., Ranjzad, P., Tengku, F., Silajdžić, E., Forster, D., Asselin, M.C., Lewis, P., Lennon, R., Plagge, A., Wang, Q., et al. (2018). Generation of Functioning Nephrons by Implanting Human Pluripotent Stem Cell-Derived Kidney Progenitors. Stem Cell Reports 10, 766–779.

van den Berg, C.W., Ritsma, L., Avramut, M.C., Wiersma, L.E., van den Berg, B.M., Leuning, D.G., Lievers, E., Koning, M., Vanslambrouck, J.M., Koster, A.J., et al. (2018). Renal Subcapsular Transplantation of PSC-Derived Kidney Organoids Induces Neo-vasculogenesis and Significant Glomerular and Tubular Maturation In Vivo. Stem Cell Reports 10, 751–765.

Boreström, C., Jonebring, A., Guo, J., Palmgren, H., Cederblad, L., Forslöw, A., Svensson, A., Söderberg, M., Reznichenko, A., Nyström, J., et al. (2018). A CRISP(e)R view on kidney organoids allows generation of an induced pluripotent stem cell–derived kidney model for drug discovery. Kidney Int.

Brockschmidt, A., Chung, B., Weber, S., Fischer, D.C., Kolatsi-Joannou, M., Christ, L., Heimbach, A., Shtiza, D., Klaus, G., Simonetti, G.D., et al. (2012). CHD1L: A new candidate gene for congenital anomalies of the kidneys and urinary tract (CAKUT). Nephrol. Dial. Transplant. 27, 2355–2364.

Brown, E.J., Pollak, M.R., and Barua, M. (2014). Genetic testing for nephrotic syndrome and FSGS in the era of next-generation sequencing. Kidney Int. 85, 1030–1038.

Burrows, C.K., Banovich, N.E., Pavlovic, B.J., Patterson, K., Gallego Romero, I., Pritchard, J.K., and Gilad, Y. (2016). Genetic Variation, Not Cell Type of Origin, Underlies the Majority of Identifiable Regulatory Differences in iPSCs. PloS Genet. 12, 1–18.

Carmosino, M., Rizzo, F., Procino, G., Basco, D., Valenti, G., Forbush, B., Schaeren-Wiemers, N., Caplan, M.J.J., and Svelto, M. (2010). MAL/VIP17, a new player in the regulation of NKCC2 in the kidney. Mol. Biol. Cell 21, 3985–3997.

Caroleo, M.C., Carito, V., Pingitore, A., Perrotta, I.D., Perri, M., Mancuso, D., Russo, A., and Cione, E. (2015). Human kidney podocyte cell population as a novel biological target of nerve growth factor. Growth Factors.

Christensen, E.I., and Birn, H. (2002). Megalin and cubilin: Multifunctional endocytic receptors. Nat. Rev. Mol. Cell Biol.

Coresh, J., Selvin, E., Stevens, L.A., Manzi, J., Kusek, J.W., Eggers, P., Van Lente, F., and Levey, A.S. (2007). Prevalence of Chronic Kidney Disease in the United States. Jama 298, 2038.

Cruz, N.M., Song, X., Czerniecki, S.M., Gulieva, R.E., Churchill, A.J., Kim, Y.K., Winston, K., Tran, L.M., Diaz, M.A., Fu, H., et al. (2017a). Organoid cystogenesis reveals a critical role of microenvironment in human polycystic kidney disease. Nat. Mater. 16, 1112–1119.

Cruz, N.M., Song, X., Czerniecki, S.M., Gulieva, R.E., Churchill, A.J., Kim, Y.K., Winston, K., Tran, L.M., Diaz, M.A., Fu, H., et al. (2017b). Organoid cystogenesis reveals a critical role of microenvironment in human polycystic kidney disease. Nat. Mater. 16, 1112–1119.

Dressler, G.R. (2006). The Cellular Basis of Kidney Development. Annu. Rev. Cell Dev. Biol.

Féraud, O., Valogne, Y., Melkus, M.W., Zhang, Y., Oudrhiri, N., Haddad, R., Daury, A., Rocher, C., Larbi, A., Duquesnoy, P., et al. (2016). Donor dependent variations in hematopoietic differentiation among embryonic and induced pluripotent stem cell lines. PloS One 11, 1–22.

Ferrara, N. (1999). Role of vascular endothelial growth factor in the regulation of angiogenesis. Kidney Int. 56, 794–814.

Fischer, E.A., Verpont, M.C., Garrett-Sinha, L.A., Ronco, P.M., and Rossert, J.A. (2001). Klf6 is a zinc finger protein expressed in a cell-specific manner during kidney development. J Am Soc Nephrol.

Forbes, T.A., Howden, S.E., Lawlor, K., Phipson, B., Maksimovic, J., Hale, L., Wilson, S., Quinlan, C., Ho, G., Holman, K., et al. (2018a). Patient-iPSC-Derived Kidney Organoids Show Functional Validation of a Ciliopathic Renal Phenotype and Reveal Underlying Pathogenetic Mechanisms. Am. J. Hum. Genet. 102, 816–831.

Forbes, T.A., Howden, S.E., Lawlor, K., Phipson, B., Maksimovic, J., Hale, L., Wilson, S., Quinlan, C., Ho, G., Holman, K., et al. (2018b). Patient-iPSC-Derived Kidney Organoids Show Functional Validation of a Ciliopathic Renal Phenotype and Reveal Underlying Pathogenetic Mechanisms. Am. J. Hum. Genet. 102, 816–831.

Freedman, B.S., Brooks, C.R., Lam, A.Q., Fu, H., Morizane, R., Agrawal, V., Saad, A.F., Li, M.K., Hughes, M.R., Werff, R. Vander, et al. (2015). Modelling kidney disease with CRISPR-mutant kidney organoids derived from human pluripotent epiblast spheroids. Nat. Commun. 6, 8715.

Graham, V., Khudyakov, J., Ellis, P., and Pevny, L. (2003). SOX2 functions to maintain neural progenitor identity. Neuron 39, 749–765.

Greka, A., and Mundel, P. (2012). Cell biology and pathology of podocytes. Annu. Rev. Physiol. 74, 299–323.

He, L., Sun, Y., Patrakka, J., Mostad, P., Norlin, J., Xiao, Z., Andrae, J., Tryggvason, K., Samuelsson, T., Betsholtz, C., et al. (2007). Glomerulus-specific mRNA transcripts and proteins identified through kidney expressed sequence tag database analysis. Kidney Int. 71, 889–900.

Hildebrandt, F. (2010). Genetic kidney diseases. Lancet 375, 1287–1295.

Iglesias, D.M., Hueber, P.-A., Chu, L., Campbell, R., Patenaude, A.-M., Dziarmaga, A.J., Quinlan, J., Mohamed, O., Dufort, D., and Goodyer, P.R. (2007). Canonical WNT signaling during kidney development. Am. J. Physiol. Physiol. 293, F494–F500.

Inrig, J.K., Califf, R.M., Tasneem, A., Vegunta, R.K., Molina, C., Stanifer, J.W., Chiswell, K., and Patel, U.D. (2014). The landscape of clinical trials in nephrology: A systematic review of clinicaltrials.gov. Am. J. Kidney Dis. 63, 771–780.

Kanno, S., Oda, N., Abe, M., Terai, Y., Ito, M., Shitara, K., Tabayashi, K., Shibuya, M., and Sato, Y. (2000). Roles of two VEGF receptors, Flt-1 and KDR, in the signal transduction of VEGF effects in human vascular endothelial cells. Oncogene 19, 2138–2146.

Kirby, A., Gnirke, A., Jaffe, D.B., Barešová, V., Pochet, N., Blumenstiel, B., Ye, C., Aird, D., Stevens, C., Robinson, J.T., et al. (2013). Mutations causing medullary cystic kidney disease type 1 lie in a large VNTR in MUC1 missed by massively parallel sequencing. Nat. Genet.

Kitamoto, Y., Tokunaga, H., and Tomita, K. (1997). Vascular endothelial growth factor is an essential molecule for mouse kidney development: Glomerulogenesis and nephrogenesis. J. Clin. Invest. 99, 2351–2357.

Kowalczyk, M.S., Tirosh, I., Heckl, D., Rao, T.N., Dixit, A., Haas, B.J., Schneider, R.K., Wagers, A.J., Ebert, B.L., and Regev, A. (2015). Single-cell RNA-seq reveals changes in cell cycle and differentiation programs upon aging of hematopoietic stem cells. Genome Res. 25, 1860–1872.

Kreidberg, J.A., Sariola, H., Loring, J.M., Maeda, M., Pelletier, J., Housman, D., and Jaenisch, R. (1993). WT-1 is required for early kidney development. Cell 74, 679–691.

LeBleu, V.S.S., Teng, Y., O’Connell, J.T.T., Charytan, D., Müller, G.A.A., Müller, C.A.A., Sugimoto, H., and Kalluri, R. (2013). Identification of human epididymis protein-4 as a fibroblast-derived mediator of fibrosis. Nat. Med. 19, 227–231.

Lee, J.W., Chou, C.-L., and Knepper, M.A. (2015). Deep Sequencing in Microdissected Renal Tubules Identifies Nephron Segment-Specific Transcriptomes. J. Am. Soc. Nephrol.

Leroy, X., Copin, M.C., Devisme, L., Buisine, M.P., Aubert, J.P., Gosselin, B., and Porchet, N. (2002). Expression of human mucin genes in normal kidney and renal cell carcinoma. Histopathology.

Li, M., Armelloni, S., Zennaro, C., Wei, C., Corbelli, A., Ikehata, M., Berra, S., Giardino, L., Mattinzoli, D., Watanabe, S., et al. (2015). BDNF repairs podocyte damage by microRNA-mediated increase of actin polymerization. J. Pathol.

Lindström, N.O., Guo, J., Kim, A.D., Tran, T., Guo, Q., De Sena Brandine, G., Ransick, A., Parvez, R.K., Thornton, M.E., Baskin, L., et al. (2018a). Conserved and Divergent Features of Mesenchymal Progenitor Cell Types within the Cortical Nephrogenic Niche of the Human and Mouse Kidney. J. Am. Soc. Nephrol. 29, 806–824.

Lindström, N.O., De Sena Brandine, G., Tran, T., Ransick, A., Suh, G., Guo, J., Kim, A.D., Parvez, R.K., Ruffins, S.W., Rutledge, E.A., et al. (2018b). Progressive Recruitment of Mesenchymal Progenitors Reveals a Time-Dependent Process of Cell Fate Acquisition in Mouse and Human Nephrogenesis. Dev. Cell 45, 651–660.e4.

Little, M.H., and McMahon, A.P. (2012). Mammalian kidney development: Principles, progress, and projections. Cold Spring Harb. Perspect. Biol. 4, 3.

Little, M.H., Combes, A.N., and Takasato, M. (2016). Understanding kidney morphogenesis to guide renal tissue regeneration. Nat. Rev. Nephrol. 12, 624–635.

Menon, R., Otto, E.A., Kokoruda, A., Zhou, J., Zhang, Z., Yoon, E., Chen, Y.-C., Troyanskaya, O., Spence, J.R., Kretzler, M., et al. (2018). Single-cell analysis of progenitor cell dynamics and lineage specification in the human fetal kidney. Development 145, dev164038.

Metsuyanim, S., Harari-Steinberg, O., Buzhor, E., Omer, D., Pode-Shakked, N., Ben-, H., Hur, Halperin, R., Schneider, D., and Dekel, B. (2009). Expression of Stem Cell Markers in the Human Fetal Kidney. PloS One 4, e6709.

Morizane, R., and Bonventre, J. V. (2017a). Generation of nephron progenitor cells and kidney organoids from human pluripotent stem cells. Nat. Protoc. 12, 195–207.

Morizane, R., and Bonventre, J. V (2017b). Generation of nephron progenitor cells and kidney organoids from human pluripotent stem cells. Nat. Protoc. 12, 195–207.

Morizane, R., Lam, A.Q., Freedman, B.S., Kishi, S., Valerius, M.T., and Bonventre, J. V. (2015). Nephron organoids derived from human pluripotent stem cells model kidney development and injury. Nat. Biotechnol. 33, 1193–1200.

Mundel, P., Heid, H.W., Mundel, T.M., Krüger, M., Reiser, J., and Kriz, W. (1997). Synaptopodin: An actin-associated protein in telencephalic dendrites and renal podocytes. J. Cell Biol. 139, 193–204.

Nguyen, Q.H., Lukowski, S.W., Chiu, H.S., Senabouth, A., Bruxner, T.J.C., Christ, A.N., Palpant, N.J., and Powell, J.E. (2010). Single-cell RNA-seq of human induced pluripotent stem cells reveals cellular heterogeneity and cell state transitions between subpopulations. 1053–1066.

Ohba, S., He, X., Hojo, H., and McMahon, A.P. (2015). Distinct Transcriptional Programs Underlie Sox9 Regulation of the Mammalian Chondrocyte. Cell Rep. 12, 229–243.

Pandey, S., Shekhar, K., Regev, A., and Schier, A.F. (2018). Comprehensive Identification and Spatial Mapping of Habenular Neuronal Types Using Single-Cell RNA-Seq. Curr. Biol. 28, 1052–1065.e7.

Pattaro, C., Teumer, A., Gorski, M., Chu, A.Y., Li, M., Mijatovic, V., Garnaas, M., Tin, A., Sorice, R., Li, Y., et al. (2016). Genetic associations at 53 loci highlight cell types and biological pathways relevant for kidney function. Nat. Commun. 7, 1–19.

Phipson, B., Er, P.X., Combes, A.N., Forbes, T.A., Howden, S.E., Zappia, L., Yen, H.-J., Lawlor, K.T., Hale, L.J., Sun, J., et al. (2019). Evaluation of variability in human kidney organoids. Nat. Methods 16, 79–87.

Piscione, T.D., Wu, M.Y.J., and Quaggin, S.E. (2004). Expression of Hairy/Enhancer of Split genes, Hes1 and Hes5, during murine nephron morphogenesis. Gene Expr. Patterns.

Poh, Y.C., Chen, J., Hong, Y., Yi, H., Zhang, S., Chen, J., Wu, D.C., Wang, L., Jia, Q., Singh, R., et al. (2014). Generation of organized germ layers from a single mouse embryonic stem cell. Nat. Commun. 5, 1–12.

Prozialeck, W.C., Lamar, P.C., and Appelt, D.M. (2004). Differential expression of E-cadherin, N-cadherin and beta-catenin in proximal and distal segments of the rat nephron. BMC Physiol.

Przepiorski, A., Sander, V., Tran, T., Hollywood, J.A., Sorrenson, B., Shih, J.-H., Wolvetang, E.J., McMahon, A.P., Holm, T.M., and Davidson, A.J. (2018). A Simple Bioreactor-Based Method to Generate Kidney Organoids from Pluripotent Stem Cells. Stem Cell Reports 11, 470–484.

Schwab, K., Patterson, L.T., Aronow, B.J., Luckas, R., Liang, H.C., and Potter, S.S. (2003). A catalogue of gene expression in the developing kidney. Kidney Int. 64, 1588–1604.

Sharmin, S., Taguchi, A., Kaku, Y., Yoshimura, Y., Ohmori, T., Sakuma, T., Mukoyama, M., Yamamoto, T., Kurihara, H., and Nishinakamura, R. (2016). Human Induced Pluripotent Stem Cell-Derived Podocytes Mature into Vascularized Glomeruli upon Experimental Transplantation. J. Am. Soc. Nephrol. 27, 1778–1791.

Stan, R. V., Tse, D., Deharvengt, S.J., Smits, N.C., Xu, Y., Luciano, M.R., McGarry, C.L., Buitendijk, M., Nemani, K. V., Elgueta, R., et al. (2012). The Diaphragms of Fenestrated Endothelia: Gatekeepers of Vascular Permeability and Blood Composition. Dev. Cell.

Taguchi, A., and Nishinakamura, R. (2017). Higher-Order Kidney Organogenesis from Pluripotent Stem Cells. Cell Stem Cell 21, 730–746.e6.

Taguchi, A., Kaku, Y., Ohmori, T., Sharmin, S., Ogawa, M., Sasaki, H., and Nishinakamura, R. (2014). Redefining the in vivo origin of metanephric nephron progenitors enables generation of complex kidney structures from pluripotent stem cells. Cell Stem Cell 14, 53–67.

Takasato, M., Er, P.X., Chiu, H.S., Maier, B., Baillie, G.J., Ferguson, C., Parton, R.G., Wolvetang, E.J., Roost, M.S., Chuva de Sousa Lopes, S.M., et al. (2015a). Kidney organoids from human iPS cells contain multiple lineages and model human nephrogenesis. Nature 526, 564–568.

Takasato, M., Er, P.X., Chiu, H.S., Maier, B., Baillie, G.J., Ferguson, C., Parton, R.G., Wolvetang, E.J., Roost, M.S., Chuva de Sousa Lopes, S.M., et al. (2015b). Kidney organoids from human iPS cells contain multiple lineages and model human nephrogenesis. Nature 526, 564–568.

Takasato, M., Er, P.X., Chiu, H.S., and Little, M.H. (2016a). Generation of kidney organoids from human pluripotent stem cells. Nat. Protoc. 11, 1681–1692.

Takasato, M., Er, P.X., Chiu, H.S., and Little, M.H. (2016b). Generation of kidney organoids from human pluripotent stem cells. Nat. Protoc. 11, 1681–1692.

Tanigawa, S., Islam, M., Sharmin, S., Naganuma, H., Yoshimura, Y., Haque, F., Era, T., Nakazato, H., Nakanishi, K., Sakuma, T., et al. (2018a). Organoids from Nephrotic Disease-Derived iPSCs Identify Impaired NEPHRIN Localization and Slit Diaphragm Formation in Kidney Podocytes. Stem Cell Reports 11.

Tanigawa, S., Islam, M., Sharmin, S., Naganuma, H., Yoshimura, Y., Haque, F., Era, T., Nakazato, H., Nakanishi, K., Sakuma, T., et al. (2018b). Renal Subcapsular Transplantation of PSC-Derived Kidney Organoids Induces Neo-vasculogenesis and Significant Glomerular and Tubular Maturation In Vivo. Stem Cell Reports 12, 751–765.

Tsankov, A.M., Akopian, V., Pop, R., Chetty, S., Gifford, C.A., Daheron, L., Tsankova, N.M., and Meissner, A. (2015). A qPCR ScoreCard quantifies the differentiation potential of human pluripotent stem cells. Nat. Biotechnol. 33, 1182–1192.

Tufro, a, Norwood, V.F., Carey, R.M., and Gomez, R. a (1999). Vascular endothelial growth factor induces nephrogenesis and vasculogenesis. J. Am. Soc. Nephrol. 10, 2125–2134.

Vivante, A., and Hildebrandt, F. (2016). Exploring the genetic basis of early-onset chronic kidney disease. Nat. Rev. Nephrol. 12, 133–146.

Wang, P., Chen, Y., Yong, J., Cui, Y., Wang, R., Wen, L., Qiao, J., and Tang, F. (2018). Dissecting the Global Dynamic Molecular Profiles of Human Fetal Kidney Development by Single-Cell RNA Sequencing. Cell Rep. 24, 3554–3567.e3.

Warejko, J.K., Tan, W., Daga, A., Schapiro, D., Lawson, J.A., Shril, S., Lovric, S., Ashraf, S., Rao, J., Hermle, T., et al. (2018). Whole exome sequencing of patients with steroid-resistant nephrotic syndrome. Clin. J. Am. Soc. Nephrol. 13, 53–62.

Wellik, D.M., Hawkes, P.J., and Capecchi, M.R. (2002). Hox11 paralogous genes are essential for metanephric kidney induction. Genes Dev. 16, 1423–1432.

Wharram, B.L., Goyal, M., Gillespie, P.J., Wiggins, J.E., Kershaw, D.B., Holzman, L.B., Dysko, R.C., Saunders, T.L., Samuelson, L.C., and Wiggins, R.C. (2000). Altered podocyte structure in GLEPP1 (Ptpro) -deficient mice associated with hypertension and low glomerular filtration rate. J. Clin. Invest. 106, 1281–1290.

Wu, H., Uchimura, K., Donnelly, E.L., Kirita, Y., Morris, S.A., and Humphreys, B.D. (2018). Comparative Analysis and Refinement of Human PSC-Derived Kidney Organoid Differentiation with Single-Cell Transcriptomics. Cell Stem Cell.

Xie, Y., Sakatsume, M., Nishi, S., Narita, I., Arakawa, M., and Gejyo, F. (2001). Expression, roles, receptors, and regulation of osteopontin in the kidney. Kidney Int.

Xinaris, C., Benedetti, V., Rizzo, P., Abbate, M., Corna, D., Azzollini, N., Conti, S., Unbekandt, M., Davies, J.A., Morigi, M., et al. (2012). In Vivo Maturation of Functional Renal Organoids Formed from Embryonic Cell Suspensions. J. Am. Soc. Nephrol. 23, 1857–1868.

Yi-Rong Peng, Karthik Shekhar, Wenjun Yan, Dustin Herrmann, Anna Sappington, Greg S. Bryman, Tavé van Zyl, Michael Tri. H. Do, A.R. and J.R.S. (2018). MOLECULAR CLASSIFICATION AND COMPARATIVE TAXONOMICS OF FOVEAL AND PERIPHERAL CELLS IN PRIMATE RETINA. Bioarxiv.

Young, M.D., Mitchell, T.J., Vieira Braga, F.A., Tran, M.G.B., Stewart, B.J., Ferdinand, J.R., Collord, G., Botting, R.A., Popescu, D.M., Loudon, K.W., et al. (2018). Single-cell transcriptomes from human kidneys reveal the cellular identity of renal tumors. Science (80-.). 361, 594–599.

Yuan, L., Le Bras, A., Sacharidou, A., Itagaki, K., Zhan, Y., Kondo, M., Carman, C. V., Davis, G.E., Aird, W.C., and Oettgen, P. (2012). ETS-related gene (ERG) controls endothelial cell permeability via transcriptional regulation of the claudin 5 (CLDN5) gene. J. Biol. Chem. 287, 6582–6591.

Živná, M., Kidd, K., Přistoupilová, A., Barešová, V., DeFelice, M., Blumenstiel, B., Harden, M., Conlon, P., Lavin, P., Connaughton, D.M., et al. (2018). Noninvasive Immunohistochemical Diagnosis and Novel *MUC1* Mutations Causing Autosomal Dominant Tubulointerstitial Kidney Disease. J. Am. Soc. Nephrol.

## Reference

Benjamini, Y., Hochberg, Y., and Benjamini, Yoav, H.Y. (1995). Benjamini and Y FDR.pdf. J. R. Stat. Soc. Ser. B 57, 289–300.

Butler, A., Hoffman, P., Smibert, P., Papalexi, E., and Satija, R. (2018). Integrating single-cell transcriptomic data across different conditions, technologies, and species. Nat. Biotechnol. 36, 411–420.

Liaw, A and Wiener, M. (2002). Classification and Regression by randomForest. R News 2(3), 18–22.

Morizane, R., and Bonventre, J. V. (2017). Generation of nephron progenitor cells and kidney organoids from human pluripotent stem cells. Nat. Protoc. 12, 195–207.

Takasato, M., Er, P.X., Chiu, H.S., and Little, M.H. (2016). Generation of kidney organoids from human pluripotent stem cells. Nat. Protoc. 11, 1681–1692.

Wickham, H. (2016). ggplot2: Elegant Graphics for Data Analysis.

