## Supplemental Tables for "Kidney organoid reproducibility across multiple human iPSC lines and diminished off target cells after transplantation revealed by single cell transcriptomics": ST1_canonical_marker_list_alldata.docx

| **Kidney compartment** |  | **Canonical Markers used for identification** | |
| --- | --- | --- | --- |
|  | **Organoid** | **Human** | **Mouse** |
| Podocyte | NPHS2, SYNPO,WT1 | NPHS2, SYNPO, WT1 |  |
| Proximal Tubule | CUBN, LRP2,AQP1 | CUBN, LRP2,AQP1 |  |
| Thick Ascending Limb | SLC12A1 | SLC12A1 |  |
| Distal Convoluted Tubule | GATA3, CDH1, MUC1 | SLC12A3, MUC1, CDH1 |  |
| Distal Nephron |  | - |  |
| Ureteric Bud |  | - |  |
| Collecting Duct – Principal Cells | - | AQP2 |  |
| Collecting Duct – alpha Intercalated Cells | - | SLC4A1 |  |
| Collecting Duct – beta Intercalated Cells | - | SLC26A4 |  |
| Mesenchymal/vascular smooth muscle | COL3A1, MEIS2, MEIS1 | ACTA2 |  |
| Mesenchymal/fibroblast |  | COL3A1 |  |
| Endothelial Cell | PECAM1 | PECAM1, KDR | Pecam1, Kdr |
| Fenestrated Endothelial Cell | - | PECAM1, PLVAP | Pecam1, Plvap |
| Immune Cells | - | PTPRC |  |

| **Other** | **Organoid** |
| --- | --- |
| Epithelial | EPCAM |
| Neuronal | STMN2, SOX11 |
| Neuronal (progenitor) | SOX2 |
| Muscle | MYOG, MYLPF |
| Melanoma-like | PMEL |

| **Other** | **Organoid** |
| --- | --- |
| Nephron Progenitor Cells (Distal) | POU3F3, PAX8 |
| Nephron Progenitor Cells (Proximal) | STMN2, SOX11 |
