## Supplemental Tables for "Kidney organoid reproducibility across multiple human iPSC lines and diminished off target cells after transplantation revealed by single cell transcriptomics": ST2_datadriven_markers.docx

| **Organoid Stage** | **Kidney Cell** | **Data-driven Marker** |
| --- | --- | --- |
| D15 | Podocyte | CLDN5 |
|  |  | SOST |
|  |  | SPARC |
|  |  | BST2 |
| D29 | Proximal Tubule (Proximal-distal gradient) | APOE |
|  | Distal Nephron | WFDC2 |
|  |  | MAL |
|  |  | DEFB1 |
